# Structural Repertoire of HCV Broadly Neutralizing Antibodies Targeting the E2 Front Layer Supersite

**DOI:** 10.1101/2025.08.04.668577

**Authors:** Xander E. Wilcox, Rajat Punia, Jessica Mimms, Nicole Frumento, Justin R. Bailey, Andrew I. Flyak

**Affiliations:** Department of Microbiology and Immunology, Cornell University, Ithaca, NY, USA; Division of Infectious Diseases, Department of Medicine, Johns Hopkins University School of Medicine, Baltimore, MD, USA; Peter Medawar Building for Pathogen Research, Nuffield Department of Medicine, University of Oxford, Oxford, UK

**Author notes:** Correspondence (A.I.F).

**Keywords:** Hepatitis C Virus, E2 glycoprotein, front layer, vaccine development

## Abstract

Structural studies of the hepatitis C virus (HCV) E2 glycoprotein in complex with broadly neutralizing antibodies (bNAbs) have been instrumental in mapping neutralizing epitopes and guiding the rational design of immunogens. However, robust structural classification of HCV bNAbs is lacking, complicating immunogen design. The majority of HCV bNAbs recognize the E2 front layer (FRLY) supersite. Here, we developed a roadmap for the structural classification of FRLY-specific bNAbs. We discovered three distinct structural classes, each utilizing a unique binding mode to engage the FRLY supersite. HCV strains with multiple FRLY polymorphisms had a profound impact on binding and neutralization of bNAbs from distinct FRLY classes. Our findings establish the FRLY as a major antigenic supersite targeted by three bNAb classes and underscore the intrinsic structural plasticity of *V_H_1-69*-encoded HCV bNAbs.

## INTRODUCTION

Hepatitis C virus (HCV) chronically infects over 50 million people worldwide, with one million new infections each year ^1^. As the leading cause of liver disease, liver cancer, and liver transplants, HCV is responsible for over 240,000 deaths globally^2^. In 2021, the World Health Organization (WHO) developed guidelines for global viral hepatitis elimination, yet the response is not on track to achieve these goals by 2030^3^. Although direct-acting antiviral (DAA) therapies to cure HCV are highly effective, several barriers, including the cost-prohibitive access to treatment^4^, the high incidence of reinfection in at-risk populations ^5^, and DAA resistance mutations^6^ indicate that control of the HCV epidemic with DAAs alone will not be possible, necessitating the development of a prophylactic vaccine.

There are several challenges to the development of a broadly protective HCV vaccine. First, HCV exhibits remarkable genetic diversity, which has been estimated to be higher than the genetic diversity of human immunodeficiency virus-1 (HIV-1)^7^. Second, the error-prone polymerase of HCV, coupled with immune pressure during natural infection, generates viral quasispecies with the capacity for viral escape^8^. Third, the HCV E1E2 surface antigen, composed of the E1 and E2 glycoproteins, is highly flexible^9^, posing a challenge for structure-based vaccine design^10^. Lastly, adaptive immune responses to HCV infection have been shown to be delayed^11^, complicating efforts to achieve rapid protection through vaccination.

Despite these challenges, approximately 25% of acutely infected individuals spontaneously clear infection^12–15^ with high antibody neutralization titers early in infection, correlating with viral clearance^16,17^. Evidence for T and B-cell memory has been demonstrated by partial protection against subsequent infections in humans and chimpanzees^18–21^. Several studies have shown that individuals who clear primary infection rapidly develop broadly neutralizing antibodies (bNAbs) that drive the virus to an unfit state, promoting clearance of HCV infection^12,22–25^. The goal of an effective HCV vaccine is to rapidly induce high titers of bNAbs observed in individuals who spontaneously clear infection. Such vaccine-induced bNAbs will need to target conserved yet structurally dynamic epitopes on the E1E2 glycoprotein.

Multiple individuals who spontaneously cleared infection develop potent bNAbs to the well- conserved E2 front layer (FRLY, also antigenic region 3, and Domain B)^35,36^ by directly inhibiting interaction with the critical host entry factor CD81^26,27^. Structural studies of E2 in complex with bNAbs have been instrumental in mapping neutralizing epitopes^14,28–35^, uncovering mechanisms of viral escape^36–42^, and guiding the rational design of E2 immunogens^43–48^. While *V_H_1-69* V-gene encoded bNAbs are observed in immune responses to influenza^49–52^, HIV-1^53,54^, and more recently, SARS-CoV-2^55^, the majority of HCV bNAbs are encoded by *V_H_1-69*^17^.

Additionally, many *V_H_1-69* HCV bNAbs that bind the E2 FRLY share a common germline- encoded CDRH3 stabilizing disulfide motif, enabling recognition of a conserved FRLY epitope^33^. Previous studies have alluded to alternative binding modes of FRLY bNAbs that contain disulfide-stabilized CDRH3^17,33,56^, yet robust structural classification of bNAbs that recognize FRLY epitopes is lacking, complicating the rational design of immunogens that can elicit FRLY-specific bNAb responses.

Here, we developed a roadmap for the structural classification of HCV bNAbs. We show that the vast majority of structurally characterized HCV *V_H_1-69* bNAbs recognize the E2 FRLY antigenic supersite. Among FRLY bNAbs, we identified three distinct structural classes (classes I-III) that do not correspond to historical epitope-based classification schemes developed to characterize E2-specific bNAbs. We further demonstrated that each FRLY bNAb class has a unique binding mode, highlighting the intrinsic structural plasticity of *V_H_1-69*-derived HCV bNAbs. This unique structural plasticity results in FRLY bNAbs using distinct mechanisms of CD81 receptor mimicry to achieve remarkable neutralization breadth and potency. Lastly, through binding and neutralization assays, we identified HCV strains that are susceptible to viral escape by each class of FRLY bNAbs. This study demonstrates the structural mechanism behind the *V_H_1-69*-driven plasticity of FRLY bNAbs and provides a rationale for the design of immunogens that stimulate the development of multiple classes of FRLY bNAbs.

## RESULTS

### HEPC108 bNAb uses a hydrophobic CDRH3 without a disulfide motif to bind a hydrophobic pocket on the E2 FRLY

By deconvoluting the neutralization patterns of antibody responses in HCV-infected individuals, we previously identified that the human antibody response to HCV is composed of several distinct neutralizing antibody types^57^, some of which have been structurally characterized^33,47,58,59^. While FRLY-specific responses have been detected in multiple HCV-infected individuals, HEPC74 and HEPC108-like bNAb responses have been detected in individuals who cleared one infection and became subsequently reinfected. Although the epitope of HEPC74 has been extensively characterized^33,47^, the binding epitope of HEPC108 is not well defined^33,37,60^. To determine the HEPC108 epitope, we solved the crystal structure of HEPC108 bound to the E2ecto domain (residues 384-646) from the 1b09 HCV strain (E2ecto_1b09_) (**Figure 1A, Table S1**). The CDRH3 of HEPC108, which adopts a straight β-hairpin conformation without a disulfide motif, makes contact with a hydrophobic pocket formed by FRLY residues W420, F442, Y443, and back layer residues P612, and Y613 (**Figure 1B** and **Figure S1A**). In addition to hydrophobic interactions with E2, the CDRH3 of HEPC108 forms hydrogen bonds with E2 residues G418, W420, L441, T444, and Y613 at the N-terminus of the α1 helix (**Figure S1B, Table S2**). Similarly to other FRLY bNAbs, HEPC108 predominantly uses V_H_ domain residues to interact with E2, burying 804 Å^2^ (76% of the total Fab buried surface area [BSA]) (**Figure 1C** and **Table S2**). Nearly 70% of buried HEPC108 V_H_ residues are hydrophobic in nature. In addition to hydrophobic contacts made by CDRH3, F29 on CDRH1 packs against a conserved W529 on the CD81 binding loop of E2. The hydrophobic CDRH2 encoded by *V_H_1-69* approaches the E2 FRLY near the C429-C503 disulfide bond and F54 packs against the hydrophobic α2 helix. While the light chain contributes minimal interactions with a total buried surface area of 265 Å (24% of the total Fab BSA), residues in CDRL1 and CDRL3 form hydrogen bonds with E2 FRLY residues with an additional two salt bridges formed between D92 of CDRHL3 and H445 and K446 of E2 (**Figure S1C, Table S2**). Compared to other FRLY bNAbs that have been structurally characterized, HEPC108 has a remarkably high 92% germline amino acid identity (**Figure 1D**) and an average CDRH3 length of 18 residues (**Figure 1E),** suggesting that traditional vaccine strategies can be used to elicit HEPC108-like bNAbs.

**Figure 1.**
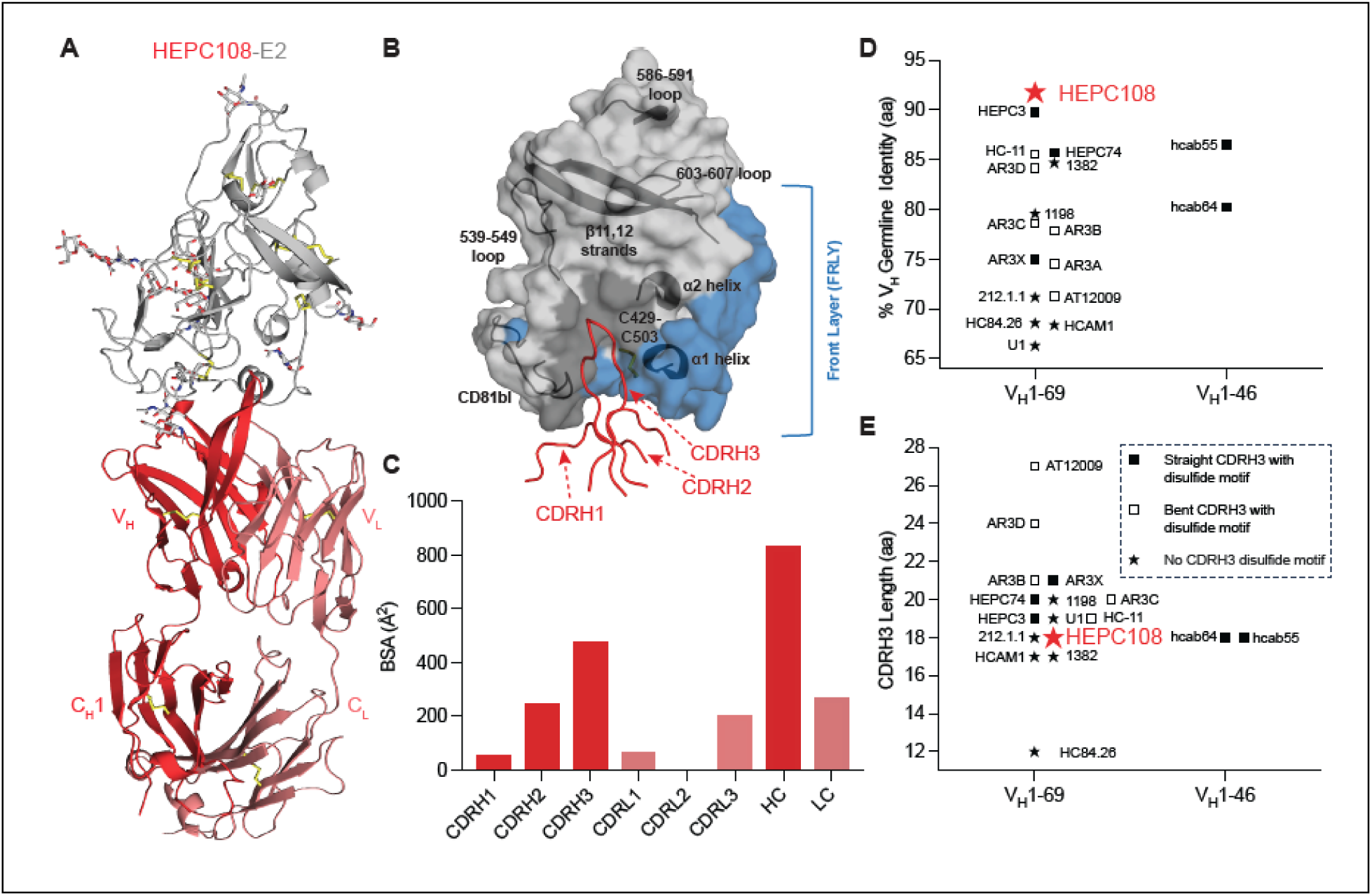
HEPC108 retains high V_H_ germline identity and targets the E2 front layer (FLRY) **(A)** Crystal structure of HEPC108-E2ecto complex. E2ecto from the 1b09 strain is shown in cartoon representation in grey with N-glycans modeled as sticks. The HEPC108 Fab is shown in a cartoon with the heavy and light chains colored in red and salmon, respectively. Disulfide bonds are shown as yellow sticks. **(B)** HEPC108 CDRH loops (colored as in **(A)**) in ribbon mapped on the surface of E2ecto with the FRLY surface colored in blue. The HEPC108 epitope is mapped to the surface of E2 in dark grey. The location of α1 and α2 helices, β11 and β12 strands, CD81 binding loop (CD81bl), loops 539-549 and 603-607 are indicated in black cartoons, and the C429-C503 disulfide bond is shown in yellow sticks. **(C)** Quantification of HEPC108 buried surface area (BSA) of E2ecto categorized by contributions from CDRs, heavy chain (HC), and light chain (LC). **(D)** V_H_-gene germline identity of known structurally characterized human FRLY-specific bNAbs. BNAbs with a straight CDRH3 containing a disulfide motif are denoted by a filled square, bNAbs with a bent CDRH3 containing a disulfide motif are denoted by an open square, and bNAbs with no CDRH3 disulfide motif are denoted by a filled star. HEPC108 bNAb is indicated by a large red star. **(E**) Distribution of CDRH3 lengths of known structurally characterized FRLY bNAbs. Coloring and symbol usage represented as in **(D)**.

### HCV bNAbs target the E2 FRLY supersite using three distinct binding modes

To see how the HEPC108 binding mode compares to other FRLY bNAbs, we aligned structures of E2 in complex with HEPC74, HEPC3, hcab55, AR3C, 1198, and 1382 Fabs. After superimposing the complexes based on their antigen structures, we quantified the relative orientations of the antibody Fab fragments by measuring the approach angle, or the minimal rotation angle needed to align one Fab to another. Approach angle (ψ) serves as a quantitative metric to characterize the E2 FRLY binding mode of HCV bNAbs. Alignment of E2-HEPC108 and E2-HEPC74 on E2 shows a rotation of ψ = 165° is required to superimpose the HEPC108 Fab onto the HEPC74 Fab (**Figure 2A, 2C**), indicating two distinct binding modes represented by HEPC74 (class I) and HEPC108 (class II). The antibody approach angles for the *V_H_1-*69- encoded HEPC3 and *V_H_1-46*-encoded hcab55 relative to HEPC74 were 15° and 25°, respectively, categorizing them as class I bNAbs (**Figure 2B, C**). Alignment of 1198 and 1382 bNAbs with HEPC108 shows these bNAbs can be classified as class II with antibody approach angles of 33° and 6° relative to HEPC108, respectively (**Figure 2B, C**). Our previous structural comparison of FRLY bNAbs with disulfide-stabilized CDRH3s indicated that the AR3C Fab was rotated by ∼77° (85° in this analysis) relative to the HEPC74 Fab, placing AR3C’s bent CDRH3 in the same region of the FRLY as HEPC74’s straight CDRH3 (**Figure 2B, 2C**)^33^, suggesting a third FRLY binding mode represented by AR3C (class III). Additional bNAbs AR3A, AR3B, and AR3D were all isolated from a single, chronically infected individual^61^ and have antibody approach angles of 11°, 18°, and 24° relative to AR3C, respectively (**Fig 2C**), indicating they belong to class III.

**Figure 2.**
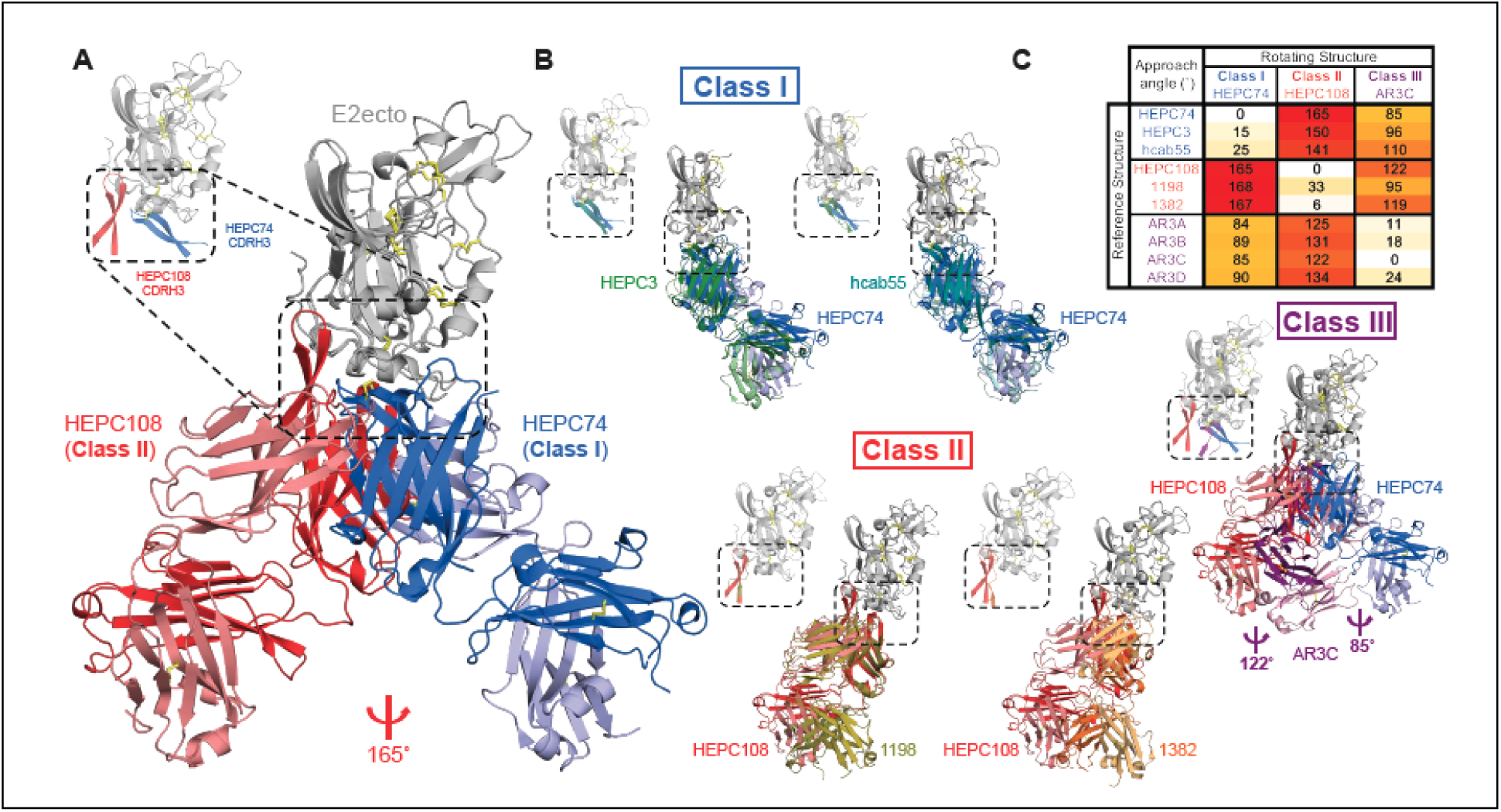
HCV E2 FRLY bNAbs can be classified into three distinct structural classes. **(A)** Structures of HEPC108 (class I bNAb) and HEPC74 (class II bNAb) in complex with E2ecto shown in cartoon aligned on the E2ecto. Coloring of HEPC108-E2ecto complex is as in Figure 1 with N-glycans omitted. HEPC74 (PDB ID: 6MEH) HC and LC are colored in dark blue and light blue, respectively. Dashed box denotes the E2 FRLY and CDR loops of the HEPC108 and HEPC74 Fabs. Left inlay displays the E2 FRLY and the Fab CDRH3 loops, highlighted with a dashed box. The red angle symbol denotes the antibody approach angle required to superimpose the HEPC108 and HEPC74 Fabs. (**B, top panel)** E2ecto in complex with E2 FRLY class I bNAbs HEPC3 (PDB ID: 6MEK) and hcab55 (PDB ID: 8W0V) aligned with HEPC74-E2ecto complex on the E2ecto. HEPC74 HC and LC colored as in **(A)** HEPC3 HC and LC colored in dark and light green, respectively. HC and LC of hcab55 are colored in dark and light teal, respectively. Disulfides are shown in yellow sticks, and dashed boxes are represented as in (**A)**. (**B, bottom panel)** E2ecto in complex with E2 FRLY class II bNAbs 1198 (PDB ID: 7RFB) and 1382 (PDB ID: 7RFC), aligned with HEPC108-E2ecto complex on the E2ecto. HEPC108 HC and LC colored as in **(A)**. 1198 HC and LC colored in deep olive and olive, respectively. 1382 HC and LC colored in orange and light orange, respectively. Disulfides are shown in yellow sticks and dashed boxes are represented as in (**A**) (**B, middle panel)** E2ecto in complex with E2 FRLY class III bNAb AR3C (PDB ID: 4MWF), aligned with HEPC108- and HEPC74-E2ecto complexes on the E2ecto. E2ecto, HEPC74 HC and LC, and HEPC108 HC and LC colored as in (**A)**. AR3C HC and LC colored in purple and light purple, respectively. Red and purple angle symbols denote antibody approach angle required to superimpose the AR3C Fab with the HEPC108 or HEPC74 Fab, respectively. Disulfides are shown in yellow sticks and dashed boxes are represented as in **(A)**. (**C)** HCV E2 FRLY-specific bNAbs segregate into three binding classes (class I-III) based on the antibody approach angle.

Mapping of representative antibodies from each class (class I: HEPC74, class II: HEPC108, class III: AR3C) onto previously defined antigenic regions, antigenic sites and antigenic domains (**Figure S2**) revealed class I-III bNAbs engage epitopes spanning antigenic region 3 (AR3^62^; residues 426-443, 529-531), antigenic site 412 (AS412^63^; residues 412-423, also referred to as domain E^64^), antigenic site 434 (AS434^65^; residues 434-446), domain D^64^ (residues 420-428, 441-443, 616), and domain B^64^ (residues 431-439, 529-535). This finding indicates that historical classification by epitope mapping does not correspond to structural classification by approach angle, which more effectively captures the structural and potentially functional differences between FRLY antibodies.

The detailed structural analysis suggests there are three distinct binding modes bNAbs use to recognize the E2 FRLY that do not correspond to historical classification schemes (**Figure S2**). Class I (HEPC74-like, including HEPC74, HEPC3, and hcab55) bNAbs place their disulfide-stabilized straight CDRH3 near the E2 C429-C503 disulfide with the glycine-rich tip of the CDRH3 packing against the W529 of the CD81 binding loop (**Figure S3, S5A**). Class II (HEPC108-like, including HEPC108, 1198, and 1382) bNAbs use their hydrophobic CDRH3 to occupy a hydrophobic pocket of E2 formed by both front and back layer residues and pack against the CD81 binding loop with the hydrophobic residues in CDRH1 (**Figure S3, S5B**).

While class III (AR3C-like, including AR3C, AR3A, AR3B, and AR3D) bNAbs bind the FRLY very similarly to class I bNAbs, antibody rotation is required for CDRH3 binding in close proximity to the C429-C503 disulfide in the FRLY^33^ (**Figure 2C**). This binding mode results in a different binding footprint of CDRH1 and CDRH2 loops, with the hydrophobic CDRH2 packing against W529 of the CD81 binding loop (**Figure S3, S5C**).

### HCV E2 antibodies cluster into six groups by inclination angle (**φ**), with FRLY antibodies further clustering into classes I-III by approach angle (**ψ**)

While the FRLY supersite is known to be the main target of bNAbs, previous studies have identified bNAbs that recognize other antigenic regions on the E1E2 heterodimer including the β-sandwich^66^, back layer^66^, and AR4/AR5^31,47,58^ antigenic regions. To unambiguously characterize the interaction modes between HCV antibodies and E2 glycoprotein and confirm the existence of three distinct binding classes among FRLY bNAbs, we performed clustering analysis using two geometric metrics: the inclination angle (φ) and the approach angle (ψ). We calculated the inclination angle (φ) between a pair of antibodies as the angle formed by lines extending from the center of mass of the E2 domain to the centers of mass of the Fv regions of antibodies (**Figure 3A**). This metric helps to distinguish antibodies targeting different antigenic regions on the E2 protein. A smaller φ indicates that antibodies target the same antigenic region, showing closely aligned orientations, while a larger φ suggests targeting of different epitopes. In contrast, the approach angle (ψ), the minimal rotation angle needed to align one Fab to another, measures variations in orientation among antibodies that bind the same antigenic region, categorizing distinct interaction modes (**Figure 3B**).

**Figure 3.**
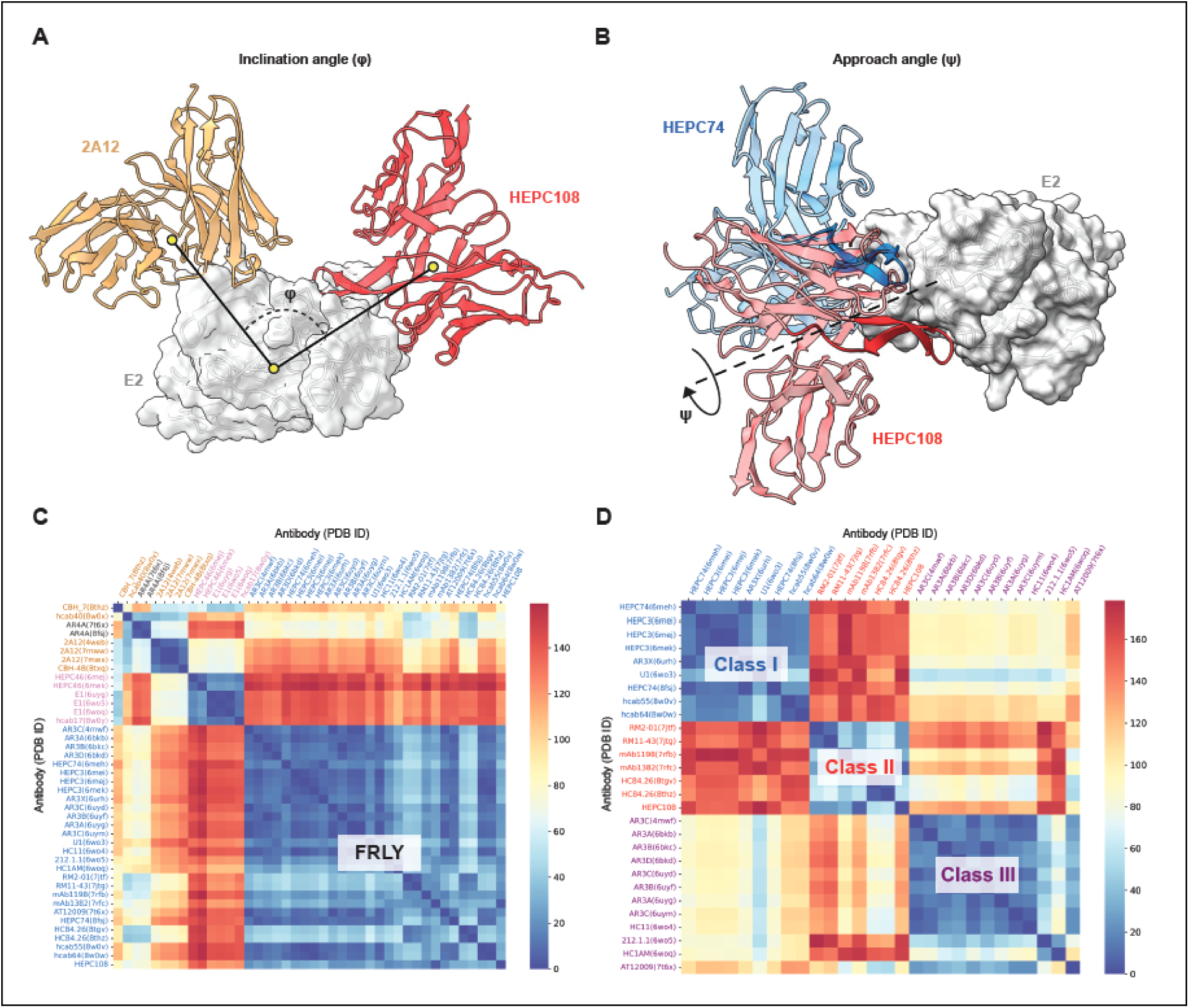
E2 FRLY supersite is targeted by bNAbs that can be clustered into three binding classes by antibody approach angle. **(A)** Schematic of inclination angle (φ) used to classify E2- specific bNAbs. Inclination angle is calculated as the angle between the center of mass of the bNAb Fab domains. E2ecto is shown in a transparent grey surface and cartoon with the HEPC108 and 2A12 Fab domains shown in red and sand cartoon, respectively. **(B)** Schematic of approach angle (ψ) used to classify E2 FRLY-specific bNAbs. The approach angle is calculated as the rotation angle required to superimpose the Fab domains after the structures are aligned on the antigen. E2ecto and HEPC108 are colored in light red and the HEPC74 Fab domain is colored in light blue, with HEPC108 and HEPC74 CDRH3 loops indicated in dark red and dark blue, respectively. **(C)** Heatmap showing inclination angle between pairs of HCV E2-binding antibodies. Clusters of antibodies with similar interaction modes were identified using the DBSCAN algorithm with cutoff angles of 25°. Axes labels, representing antibody names and PDB IDs are color-coded by cluster group where E2 FRLY antibodies are colored blue, E2 β-sandwich antibodies colored in magenta, E2 back layer antibodies colored in sand, and AR4-like antibodies colored in grey. **(D)** Heatmap showing clustering of E2 FRLY bNAbs by antibody approach angle. Clusters of antibodies with similar interaction modes were identified using the DBSCAN algorithm with cutoff angles of 45°. Axes labels, representing antibody names and PDB IDs, are color-coded by cluster group where FRLY class I bNAbs are colored in blue, class II bNAbs are colored in red, and class III bNAbs are colored in purple.

Using structural data from the Protein Data Bank, we calculated pairwise inclination angles for all available E2-antibody complexes. The DBSCAN clustering algorithm was used with a cutoff angle of 25° for φ, chosen based on the minima in the distribution histogram of pairwise angles, making the categorization minimally sensitive to the choice of cutoff (**Figure S4A**). For ψ, we selected a cutoff of 45° to classify interaction modes among antibodies targeting the same epitope (**Figure S4B**). The clustering based on φ organized the antibodies into six distinct clusters representing unique antigenic regions including the E2 FRLY, β-sandwich, back layer, and E1E2 AR4 (**Figure 3C**), with a majority of the structures (28 out of 42) targeting the FRLY antigenic supersite.

Given our previous structural analysis identifying three FRLY supersite bNAb classes, we then further clustered antibodies targeting the E2 FRLY supersite based on approach angle (ψ) and found the same three bNAb classes (**Figure 3D**). In addition to HEPC74, HEPC3, and hcab55, the clustering analysis assigned additional bNAbs AR3X, hcab64, and U1 to class I (**Figure 3D**). Our previous structural characterization of AR3X, which was isolated from the same chronically infected individual as class III bNAbs AR3A-D^61^, revealed that it engages the FRLY with a straight disulfide-stabilized CDRH3 similar to other class I bNAbs HEPC74 and HEPC3 while additionally employing an unusually long CDRH2 insertion (**Figure S5A**) to form hydrogen bonds with the FRLY. Similar to hcab55, hcab64 is another *V_H_1-46*-encoded bNAb that was isolated from an elite neutralizer who cleared three HCV infections^66^. While class I predominantly comprises bNAbs containing a disulfide-stabilized CDRH3, U1 lacks a CDRH3 disulfide-motif (**Figure S5A**) yet its CDRH3 packs against the CD81 binding loop and N-terminus of the FRLY, similar to all other class I bNAbs.

In addition to human bNAb HC84.26, clustering analyses categorized two bNAbs isolated from *Rhesus macaques* immunized with the Chiron recombinant E1E2 complex^56,67^ as class II (**Fig 3D, Figure S6**), indicating that E1E2 immunization can elicit bNAbs that resemble those isolated from infected humans but do not represent the full structural diversity of bNAbs targeting the FRLY supersite. In addition to AR3A-D bNAbs, clustering analyses assigned HC11, 212.1.1, HC1AM, and AT12009 to class III (**Figure 3D**). While AR3B, AR3D, and HC1AM lack disulfide-stabilized CDRH3s (**Figure S5C)**, the CDRH binding footprints are remarkably similar to AR3A and AR3C.

Similarly, bNAb 212.1.1 also lacks a disulfide-stabilized CDRH3 and has a similar binding footprint to other class III bNAbs. However, in this 212.1.1-E2 structure, E2 is observed in the alternate conformation (conformation B), where the α1-helix is displaced by 10 Å^68^. To date, all other E2-bNAb and E1E2-bNAb complex structures capture E2 in the A conformation, including HEPC108 which utilizes its CDRH3 to interact with back layer residue Y613 (**Figure 4A**). In contrast, bNAb 212.1.1 contacts back layer residues Y613 and W616 using the hydrophobic CDRH2 (**Figure S4C**), an interaction likely requiring α1-helix displacement (**Figure 4B**). To determine the E2 states where back layer residues Y613 and W616 are accessible for bNAb engagement, we performed 10 μs molecular dynamics (MD) simulations. In the apo (antibody- unbound) state, residues Y613 and W616 of the E2 back-layer are transiently accessible but remain predominantly buried (**Figure 4C**). Interestingly, the solvent exposure of W616 does not correlate with that of Y613, suggesting independent mechanisms of their transitions from buried to exposed states. To determine if conformation B of the E2 FRLY is sampled during MD simulations, we introduced an order parameter θ_*α*1-helix_ that effectively distinguishes conformation B (θ_*α*1-helix_ ≅ 35°) from conformation A (θ_*α*1-helix_ ≅ 90°) (**Figure 4D**). As shown in **Figure 4E**, E2 remained in conformation A throughout the simulation trajectory and no transition to conformation B was observed. This observation suggests two primary possibilities: (a) conformation B may be an induced state exclusively due to the interactions with antibody 212.1.1, or (b) the timescale of our MD simulations (∼10 μs) may be too small to capture the transition between conformations A and B. To address the latter possibility, we employed well- tempered metadynamics (wt-metad) simulations^69^, an enhanced sampling technique that introduces a bias potential along slow degrees of freedom, θ_*α*1-helix_, to overcome the kinetic barriers. The free energy landscape computed along θ_*α*1-helix_, revealed a metastable conformation near θ_*α*1-helix_ = 40°, separated from conformation A by an energy barrier of 7.2 kcal/mol (**Figure 4f**). This analysis suggests that conformation B is a metastable state of the E2 in unliganded state, although less stable compared to conformation A. E2 exist primarily in the A conformation that is recognized by all three classes of bNAbs targeting the E2 FRLY supersite.

**Figure 4.**
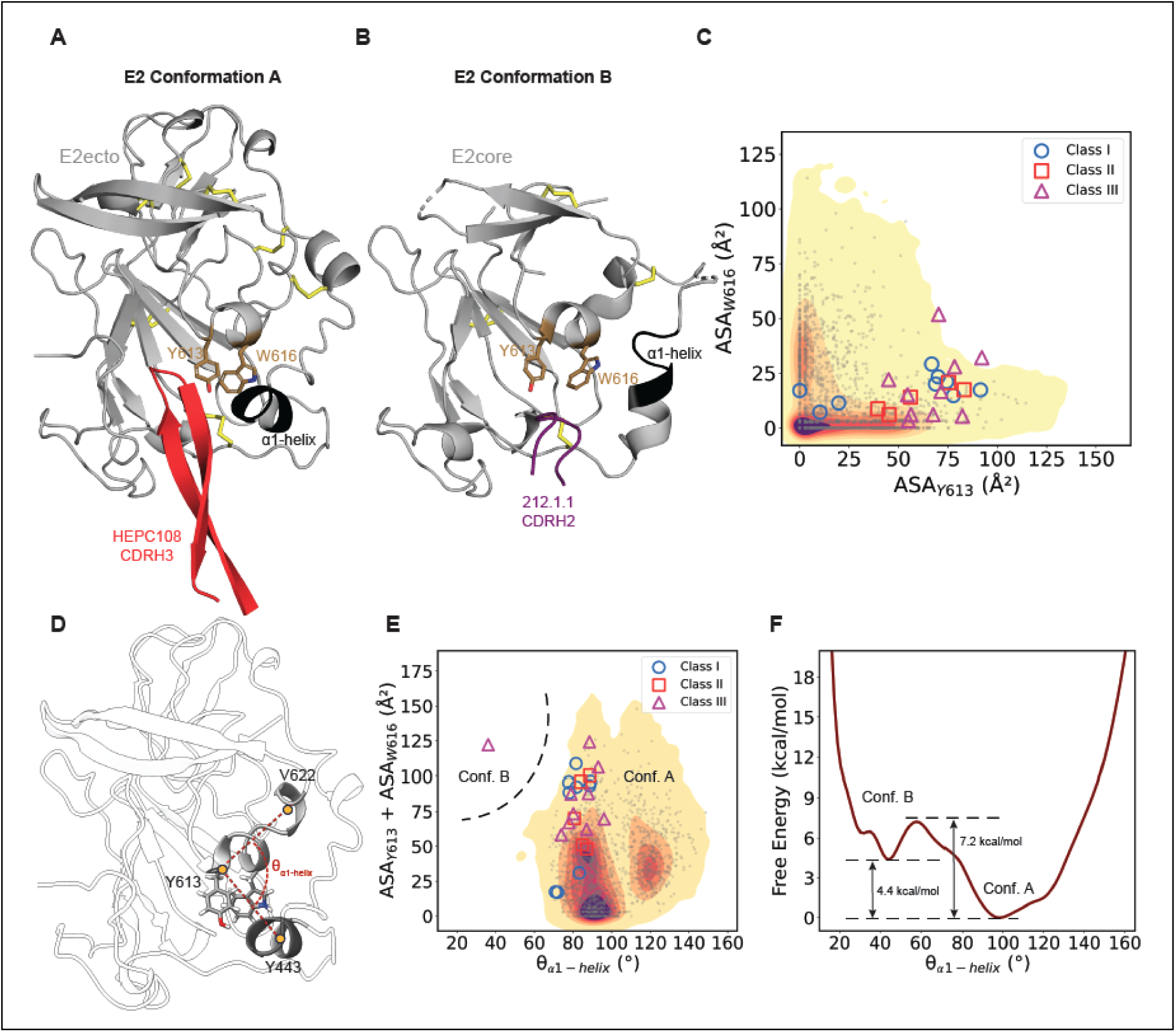
HCV E2 predominantly adopts a single conformation recognized by all three classes of FRLY bNAbs. **(A)** Structure of E2ecto-HEPC108 showing the CDRH3 loop of class II bNAb HEPC108, colored in red cartoon, interacting with back layer residue Y613 when E2ecto is in the A conformation. E2 is shown in grey cartoon with disulfides shown in yellow sticks and α1-helix colored in black. **(B)** Structure of E2core-212.1.1 complex (PDB ID: 6WO5) with E2 in the B conformation allowing CDRH2 of class II bNAb 212.1.1, colored in purple cartoon to contact back layer residues Y613 and W616. **(C)** 2D density plot of the accessible surface areas (ASA) of HCV E2 residues W616 and Y613, computed from 10 µs MD simulations. Grey dots represent MD data sampled every 5 ns. Markers indicate ASAs from E2 crystal structures in complex with various classes of FRLY antibodies, where ASA calculations included exposure to both solvent and antibody. **(D)** Order parameter θ_*α*1-helix_ defined as the angle between C*α* atoms of residues 443, 613 and 622 to differentiate E2 conformations A and B. **(E)** 2D density plot of total ASA of residues W616 and Y613, and order parameter computed from 10 µs long MD simulations. Marker notations same as **(C)**. **(F)** Free energy profile along θ_*α*1-helix_, estimated from a 500 ns long well-tempered metadynamics simulations, suggesting an existence of metastable conformation B of E2 in its apo (antibody-unbound) state.

### bNAbs converge on a single site of vulnerability in the E2 FRLY, blocking E2 binding to CD81

While at least four attachment factors have been identified to be important in HCV cellular entry, blocking E1E2 interaction with CD81 on hepatocytes completely prevents viral entry^27^. As a result, many potent bNAbs target conformational epitopes within the E2 FRLY and CD81 binding loop^28^. Previous studies have demonstrated that while E2 substitutions in the FRLY arise throughout infection, FRLY bNAbs select for escape mutations in the FRLY that drive the virus to an unfit state, mediating spontaneous clearance^24,41^. Structural analysis of E2 bound to the CD81 large extracellular loop (LEL) from tamarin shows significant overlap of the CD81 binding footprint with potent bNAbs, suggesting that direct competition for CD81 binding is the mechanism of neutralization by FRLY bNAbs (**Figure 5A**)^70^. To determine how FRLY bNAbs from each class recognize E2 and block the CD81 LEL binding site, structures of E2 bound to class I-III bNAbs were aligned with the E2-tamarin CD81 structure on the antigen. Class I bNAbs predominantly use their straight, disulfide-stabilized CDRH3 to mimic helix C of CD81 LEL with additional contribution of CDRH1 to the binding footprint, resulting in an E2 BSA of 1058 Å^2^ (**Figure 5B**). Conversely, class II bNAbs use CDRH1 to occupy the CD81 LEL helix C binding site and CDRH3 (lacking a disulfide motif) to mimic helix D of CD81 LEL, creating an overall buried E2 surface area of 967 Å^2^. Similar to class I, class III bNAbs use their bent, disulfide-stabilized CDRH3 to mimic CD81 LEL helix C with an overall total E2 buried surface of 841 Å^2^ (**Figure 5B**). Interestingly, while the CDRH2 loops of all three classes of bNAbs are predominantly encoded by the same V-gene, each binds to a different region of the E2 FRLY, further highlighting the structural plasticity of *V_H_1-69*-encoded CDRH2 loops to recognize the flexible FRLY by mimicking CD81 LEL^9,33,34,59^

**Figure 5.**
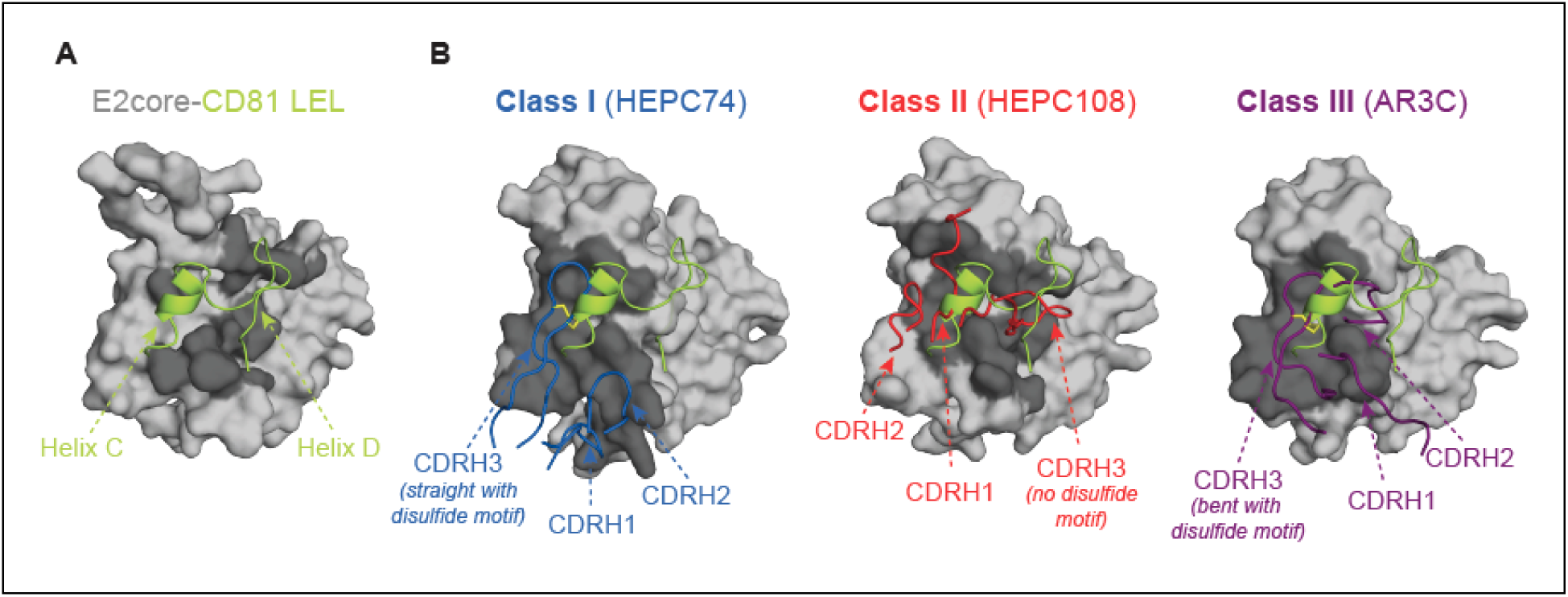
E2 FRLY supersite bNAbs mimic CD81 LEL helices C and D. **(A)** Structure of E2core in complex with tamarin CD81 LEL (PDB ID: 7MWX). CD81 LEL helices C and D are shown in chartreuse cartoon, and E2core is displayed in grey surface with the binding footprint of CD81 LEL in dark grey. **(B)** Overlay of CD81 LEL C and D helices and CDRH loops of E2 FRLY-specific HEPC74 (class I), HEPC108 (class II), and AR3C (class III) with E2ecto (HEPC108 and HEPC74 structures) or E2core (AR3C structure) rendered in grey surface with epitopes colored in dark grey. Coloring of CDRH loops as in Figure 2.

### FRLY bNAbs are susceptible to E2 polymorphisms

In order to develop a broadly protective HCV vaccine, bNAbs elicited by an immunogen must retain neutralization capacity to diverse polymorphisms existing in the viral population, as well as those that emerge throughout infection due to immune pressure. While the E2 FRLY is relatively conserved^28^, several amino acid substitutions have been identified to confer resistance to FRLY bNAbs^71,72^. Given the overlapping, but distinct binding footprints observed in each class of FRLY bNAbs, we hypothesized that amino acid substitutions in the FRLY may impact each class differently. To determine which residues were crucial for class I-III bNAb binding and thus potentially susceptible to viral escape polymorphisms, we conducted MD simulations of E2 in complex with a representative bNAb from each class. Residue-wise binding energy decomposition analyses indicated distinct E2 residues critical for each class of FRLY bNAb (**Figure 6A**). For all three classes, FRLY residues 442 and 443, as well as residue 529 in the CD81 binding loop, were found to be critical for bNAb binding. For class I bNAbs (represented by HEPC74), E2 residues 434, 438, 441, 446-448 significantly contribute to binding with residues 447 and 448 unique to class I only (**Figure 6A, Figure S7A**). While residue-wise energy binding energy decomposition analysis did not identify residue 431 as strongly contributing to the binding energy of HEPC74, the HEPC74 Fab buries 98% of the accessible surface area at this position, indicating that it likely plays a role in maintaining the E2-Fab interface. The binding footprint of class II bNAbs (represented by HEPC108) is markedly distinct, involving residues that are non-essential for class I or III binding including 420, 422, 612, and 613 (**Figure 6A, Figure S7B**). Class III bNAbs (represented by AR3C) bind E2 residues also important for class I binding including 434 and 446, with residues 427, 429, and 433 unique to the AR3C (**Figure 6A, Figure S7C**). Overall, the individual binding footprints suggests each FRLY class may be sensitive to different circulating viral polymorphisms.

**Figure 6.**
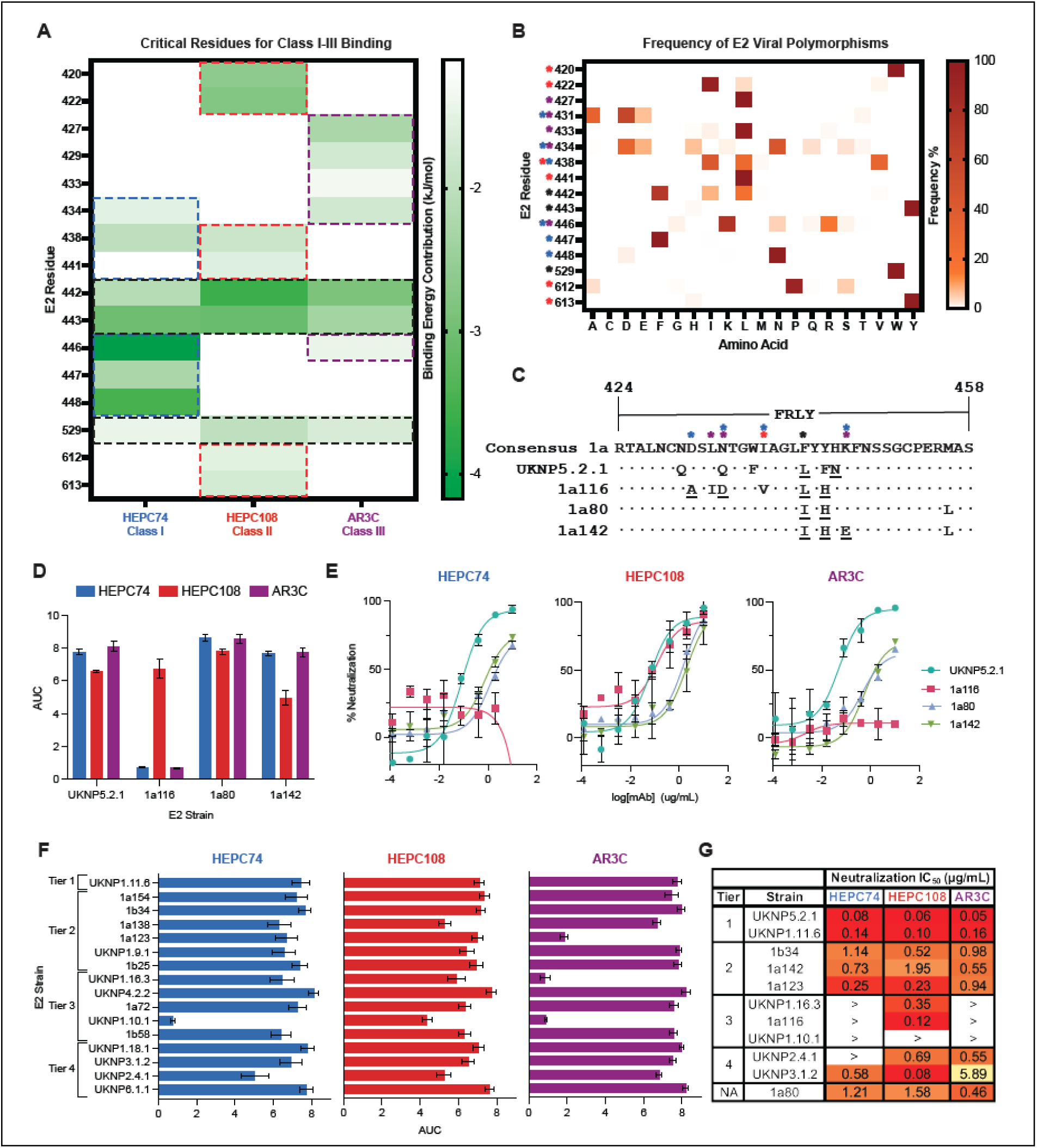
Class I-III bNAbs are susceptible to distinct E2 FRLY polymorphisms. **(A)** Heatmap of the predicted binding energy contribution of key E2 residues for HEPC74 (class I), HEPC108 (class II), and AR3C (class III) binding as determined by residue-wise energy decomposition from 10 µs long MD simulations. Residues with a binding energy contribution less than -1.3 kJ/mol were considered to significantly contribute to Fab binding. Dashed boxes indicate residues that contribute significantly to a single FRLY bNAb class with class I in blue, class II in red, and class III in purple. A black dashed box indicates residues contributing to binding of all three FRLY classes. **(B)** Frequency of E2 polymorphisms in the E2 FRLY, CD81bl, and E2 back layer. E2 residues important for class I-III binding are denoted by blue, red, and purple asterisks, and residues important for all three classes are denoted by a black asterisk. **(C)** Amino acid sequence alignment of E2 strains 1a116, 1a80, 1a142, and 1a157 compared to the genotype 1a consensus sequence. Polymorphic positions deemed to be important for class I, II and III binding determined in (**A)** are denoted by a blue, red, and purple asterisk, respectively, whereas residues important for binding by all three classes is denoted by a black asterisk. **(D)** ELISA binding data plotted as area under the curve (AUC) of HEPC108, HEPC74, and AR3C binding to E2 strains UKNP5.2.1, 1a116, 1a80, and 1a142. Error bars represent SD of three independent experiments. **(E)** Neutralization potency plots of HEPC108, HEPC74, and AR3C against UKNP5.1.2, 1a116, 1a80, 1a142 HCVpp. Error bars represent SD of duplicate measurements. **(F)** ELISA data plotted as AUC showing the binding of HEPC108, HEPC74, and AR3C to E2 strains derived from the HCVpp neutralization panel. **(G)** Summary table of neutralization potency of HEPC74 (class I), HEPC108 (class II), and AR3C (class III) bNAbs each measured against UKNP5.2.1, 1a116, 1a80, 1a142 and an abbreviated panel of 7 HCVpp. Antigens are categorized by tier, when applicable; 1a80 is not assigned a tier. The IC_50_ values for each E2-bNAb combination are shown, and the > symbol indicates IC_50_ values in which percent neutralization at the highest antibody concentration tested was lower than 50% or the IC_50_ exceeded the maximum mAb concentration tested.

To uncover specific E2 FRLY polymorphisms that may impact binding and neutralization capacity of class I-III bNAbs, we assessed the global prevalence of substitutions in the E2 FRLY by analyzing genotype 1a sequences from the HCV-Genes Linked by Underlying Evolution (GLUE) database^73,74^. Amongst the residues identified to be important for class I-III binding, we identified notable polymorphisms at E2 FRLY positions 431, 434, 438, 442, and 446 (**Figure 6B**). When comparing the critical residues for binding and the frequency of E2 FRLY polymorphisms, we hypothesized that class I and III bNAbs would be susceptible to polymorphisms at positions 431, 434 and 446 (**Figure 6A)**. In contrast to class I and III bNAbs, E2 residues critical for binding to class II bNAbs are highly conserved including frequencies in the viral population of 99.7% for W420, 90.3% for P612, and 98.7% for Y613 (**Figure 6B**) suggesting that class II bNAbs may be susceptible to neutralization escape by E2 viral polymorphisms. Two of the three residues critical for binding to E2 by all three classes are highly conserved, including Y443 in the FRLY (99.0% frequency in the viral population) and W529 in the CD81 binding loop (99.9% frequency in the viral population). Interestingly, while position 442 is critical for binding of all three classes of bNAbs, it is predominantly encoded by the bulky, hydrophobic amino acid phenylalanine (71.7% frequency), although viral polymorphisms encoding small hydrophobic residues are observed at lower frequencies (L, 19.2% and I, 8.00%). Given that F442 has been identified as a critical component of the E2 hydrophobic pocket engaged by CDRH3 loops of class II bNAbs, it is possible that variation at this position to encode smaller hydrophobic residues could impact class II bNAbs more than class I or III. Considering the distinct binding footprints of class I-III bNAbs and the presence of polymorphisms at key interaction residues, we hypothesized each class would display a unique pattern of binding and neutralization potency to strains carrying one or more FRLY polymorphisms.

### Impact of E2 polymorphisms on FRLY bNAb binding and neutralization sensitivity

To explore how FRLY E2 polymorphisms impact bNAb binding and neutralization, we selected genotype 1a strains with one or more polymorphisms that would potentially have a deleterious effect on binding of each FRLY bNAb class. The HCV strain 1a116 has been previously shown to be resistant to neutralization by class I bNAbs HEPC74 and HEPC3 and only weakly neutralized by class III bNAb AR3C, yet was strongly neutralized by class II bNAb HEPC108^33,75^. This strain harbors D431A, L433I, N434D, I438V, and F442L polymorphisms in the FRLY (**Figure 6C**), many of which were shown by MD simulations (**Figure 6A, Figure S7B**) to be important for class I-III bNAb binding. In addition, we selected 1a142, which harbors F442I and K446E polymorphisms, both implicated in class I and III bNAb binding. Finally, we included 1a80, which carries only the F442I substitution important for all three bNAb classes (**Figure 6C**). ELISA binding assays of representative class I-III bNAbs to the selected E2 strains indicated that while class II bNAb HEPC108 retained strong binding to 1a116, minimal binding of class I and III bNAbs HEPC74 and AR3C was detected (**Figure 6D, Figure S8A**). Furthermore, while all three bNAb classes bound the 1a80 and 1a142 strains, class II bNAb HEPC108 displayed approximately 1.5-fold weaker binding to 1a142 when compared to class I bNAb HEPC74 and class III bNAb AR3C (**Figure 6D, Figure S8A**). These findings highlight the differential binding patterns of class I-III bNAbs to E2 strains with specific FRLY polymorphisms, prompting further investigation into how effectively each FRLY bNAb class neutralizes these strains.

To examine how FRLY polymorphisms impact the neutralization potency of class I-III bNAbs, a representative bNAb from each class was tested against the highly neutralization-sensitive antigen UKNP5.2.1, along with 1a116, 1a80, and 1a142. All three bNAbs effectively neutralized UKNP5.2.1 (IC_50_ < 0.1 µg/mL) (**Figure 6E, 6G**). Consistent with binding studies, neutralization potency experiments showed that HEPC108 was able to efficiently neutralize 1a116 (IC_50_ = 0.12 µg/mL), whereas HEPC74 and AR3C failed to neutralize this strain (**Figure 6E, 6G**), indicating class I and III are particularly sensitive to FRLY polymorphisms present in the 1a116 strain.

Although polymorphisms in 1a116 occur at positions predicted to be important for binding of all three bNAb classes, the I438V (class I and III) and F442L (class I, II, and III) preserve the hydrophobic nature of these residues, whereas the D431A (class I) and N434D (class I and III) polymorphisms significantly alter the biochemical properties at these positions and likely negatively impact binding. Both strains 1a80 and 1a142 were more neutralization resistant by all three classes of FRLY bNAbs when compared to UKNP5.2.1, but appeared to be the most resistant to class II bNAb HEPC108 (**Figure 6E, 6G**). Overall, the neutralization data indicate that D431A and N434D FRLY polymorphisms in 1a116 have a profound impact on neutralization by class I and III bNAbs, whereas the K446E polymorphisms of 1a142 and F4421 of both 1a80 and 1a142 had minimal impact on the neutralization potency of bNAbs from all three classes.

To extend on this analysis and broadly assess the susceptibility of class I-III FRLY bNAbs to circulating viral species, the binding of a representative bNAb from each class was assayed against sixteen E2 antigens from an antigenically diverse HCV pseudoparticle panel representing HCV genotypes 1-6, tiered by increased neutralization resistance^76^ (**Figure 6F, S8B**). While class II bNAbs bound all seven neutralization-sensitive tier 1 and 2 antigens, class III bNAbs failed to bind tier 2 antigen 1a123. Binding to the entire panel of tier 3 and 4 antigens was only achieved by class II bNAb HEPC108, with class I and III bNAbs lacking binding to UKNP1.10.1 and class III bNAb AR3C additionally failing to bind UKNP1.16.3.

To assess if the observed loss of binding of class I and III bNAbs impacted neutralization, we tested on representative class I-III bNAbs against selected tier 1-4 strains (**Figure 6G, Figure S9**). All three bNAbs representing class I-III potently neutralized strains from tier 1 (UKNP1.11.6) and tier 2 (1b34 and 1a123). Class I (HEPC74) and class III (AR3C), but not class II (HEPC108) failed to neutralize all tested tier 3 strains: UKNP1.16.3, with a F442I polymorphism, 1a116 with D431A, N434D, F442L, and UKNP1.10.1, bearing D431A and N434D polymorphisms. The functional difference between class I (HEPC74) and class III (AR3C) is evident in neutralization potency against tier 4 resistant strains. While both HEPC108 and AR3C neutralize UKNP2.4.1, HEPC74 does not. Given that position 446 contributes more significantly to the binding energy of HEPC74 than AR3C (**Figure 6A**), it is likely that the K446N polymorphism, along with other factors such as FRLY dynamics, contributes to UKNP2.4.1’s resistance to HEPC74. In contrast, HEPC74 exhibits a 10-fold higher neutralization potency against another tier 4 strain, UKNP3.1.2, which lacks significant polymorphisms within the binding footprint of class I-III bNAbs, highlighting the functional differences between class I and III, as well as additional factors beyond FRLY polymorphisms that contribute to neutralization resistance. Taken together, these data demonstrate that FRLY polymorphic hotspots must be carefully considered when designing E2 immunogens displaying FRLY neutralizing epitopes.

## DISCUSSION

The majority of bNAbs isolated from HCV-infected individuals target the E2 glycoprotein FRLY, an antigenic region that interacts with the host entry factor CD81. FRLY bNAbs have been previously described based on epitope mapping as recognizing antigenic region 3 (AR3^62^; residues 426-443, 529-531). However, the binding footprint of antigenic region 3 antibodies also overlaps with several previously described antigenic sites and antigenic domains, including antigenic site 412 (AS412^63^; residues 412-423, also referred to as domain E^64^ or epitope I^77^), antigenic site 434 (AS434^65^; residues 434-446, also referred to as domain D^64^ or epitope II^77^), and domain B^64^ (residues 431-439, 529-535). Despite the existence of multiple frameworks used historically to describe FRLY antibodies, the subtle differences in antibody approach angle and binding footprints result in functional differences not captured by existing classification schemes. (**Figure S2)**. In this study, we developed a comprehensive roadmap for structural classification of FRLY bNAbs using antibody approach angle clustering and detailed structural analysis, identifying three distinct binding classes of bNAbs targeting the E2 FRLY antigenic supersite (**Figure 2, 3**) with functionally different binding and neutralization properties (**Figure 6**). FRLY bNAbs from three binding classes predominately utilize the *V_H_1-69* gene segment and unique CDRH3 properties to target overlapping but distinct FRLY epitopes, highlighting the exceptional structural plasticity of *V_H_1-69*-encoded bNAbs.

Class I and III bNAbs include antibodies previously described as recognizing AR3 or domain B. Here we determined these bNAbs can be split into two distinct classes. Class I FRLY bNAbs (HEPC74, HEPC3, AR3X, U1, hcab55, and hcab64), predominantly use a straight CDRH3 loop stabilized with a disulfide motif (**Figure 5**). Although U1 lacks a disulfide motif, the tip of its CDRH3 loop interacts with the FRLY and CD81 binding loop in a manner similar to other class I bNAbs. Class I bNAbs include reference bNAbs, such as HEPC74, HEPC3, hcab55, and hcab64, isolated from individuals who cleared one or more HCV infections^66,78^, raising the question if the emergence of FRLY class I bNAbs is associated with HCV clearance. Class III FRLY bNAbs (AR3C, AR3A, AR3B, AR3D, HC11, 212.1.1, HC1AM, and AT12009) predominantly use a bent CDRH3 loop bearing a disulfide motif, which is thought to stabilize the β-hairpin conformation and enable efficient recognition of the E2 FRLY^33^. The conformation of the bent CDRH3 results in different CDRH1 and CDHR2 binding footprints when compared to class I, resulting in different susceptibility to FRLY polymorphisms. There are exceptions of class III bNAbs with bent CDRH3s lacking a disulfide motif, including AR3B, AR3D, 212.1.1, and HC1AM, suggesting a stable CDRH3 β-hairpin may not require a disulfide motif. While many class III bNAbs have been isolated from chronically infected individuals (AR3A-D^61^and HC11), class III bNAb 212.1.1, was isolated from an individual who cleared three infections, yet this individual had the interferon-lambda beta subunit (*IL28B*) polymorphism^79^ associated with spontaneous viral clearance^80^, making it unclear if the class III bNAbs were responsible for viral clearance in this individual. Additional studies will be needed to investigate the relationship between the specificity and magnitude of class I and III FRLY responses with the HCV clearance.

Class II FRLY bNAbs (HEPC108, 1198, 1382, HC84.26, RM2-01, and RM11-43) were previously described to have a distinct binding mode from HEPC74 (class I) and AR3C (class III) like antibodies. Class II FRLY bNAbs lack a disulfide motif stabilizing their CDRH3 and use their CDRH3 (HEPC108, 1198, 1382, RM2-01, and RM11-43) to engage a hydrophobic pocket comprised of both FRLY and back layer residues. In contrast, HC84.26 uses its CDRH1 residues to bind the same hydrophobic pocket. HC84.26 was previously described as an AS434/domain D antibody. However, detailed structural analysis indicates that the HC84.26 epitope overlaps with multiple antigenic regions and sites (**Figure S2**). Interestingly, when using a more stringent cutoff angle for DBSCN clustering, HC84.26 can be separated from other class II bNAbs (**Figure S4B, S4C**). Additional structural studies will help to determine if the HC84 family of bNAbs^81^ forms a distinct class II subclass.

While all class II human bNAbs (HEPC108^75^, 1198 ^17^, 1382^17^, and HC84.26^82^) were isolated from chronically infected individuals, HEPC108 was isolated early during a second infection after the clearance of a prior infection^75^. Notably 1198 and 1382 were isolated from elite neutralizers who exhibited high neutralization titers and exceptionally broad and potent bNAbs^17^, emphasizing the need for further studies investigating the kinetics of class II FRLY bNAb responses in viral clearance and persistence. Intriguingly, class II includes bNAbs isolated from *Rhesus macaque* immunized with the Chiron E1E2 vaccine^56,67^ (**Figure 3, Figure S6**), indicating that class II bNAbs can be effectively elicited through vaccination.

While recognition of the E2 FRLY by a hydrophobic CDRH2 has been considered to be a defining feature by *V_H_1-69* bNAbs^59^, our findings show that CDRH2 loops of *V_H_1-69*-derived bNAbs are not always hydrophobic. Class I and III bNAbs have hydrophobic CDRH2 loops with 3/4 human class II and 6/8 class III bNAbs bearing F54 derived from *V_H_1-69* F alleles suggested to be important for binding and neutralization^59^. In contrast, *V_H_1-*69-derived class I bNAbs hydrophobicity scores range from -0.37 to 1.58 with only half bearing F54. Furthermore, *V_H_1-46* derived hcab55 and hcab64 have quite hydrophilic CDRH2s (hydrophobicity indexes of -0.51 and -1.12, respectively), further highlighting the significant structural plasticity of FRLY bNAbs derived from different V-genes. Additionally, our findings show that the CDRH2 loops of each class interact with different hydrophobic patches of the E2 FRLY. This exceptionally structural plasticity allows them to use distinct mechanisms of CD81 mimicry to block HCV entry into the host cells. While class I and III bNAbs predominantly use their disulfide stabilized CDRH3 loop to mimic helix C of CD81 LEL, class II bNAbs use CDRH1 to mimic helix C and CDRH3 to mimic helix D of CD81 LEL (**Figure 5**).

Given that E2 also displays high conformational dynamics, we sought to determine which E2 FRLY conformation is targeted by structurally characterized FRLY bNAbs. We found that the vast majority of FRLY bNAbs recognize E2 in the A conformation, in which α1-helix packs against hydrophobic back layer residues. The existence of an alternative E2 conformation has been recently reported: a truncated E2 from the H77 (1a154) strain, when bound by class III bNAb 212.1.1, was found to adopt the B conformation, in which α1-helix is displaced and back layer residues become accessible^68^. Through enhanced molecular dynamic simulations, we observed that the B conformation appears to exist as a metastable state and E2 predominately occupies the A conformation, which is recognized by all three FRLY supersite classes (**Figure 5**).

Lastly, we identified known E2 polymorphisms that may contribute to resistance to each class of FRLY bNAbs (**Fig 6**). We show that FRLY polymorphisms in the resistant HCV strain 1a116 prevent the binding and neutralization of representative members of class I and III bNAbs (**Figure 6C-E, 6G**). We extended this analysis to probe the binding of each FRLY supersite bNAb class to a panel of genetically and antigenically diverse E2 strains. While representative class II bNAb HEPC108 bound all 16 strains, class I representative bNAb HEPC74 and class III representative bNAb AR3C failed to bind tier 3 antigen UKNP1.10.1 with AR3C also additionally showing minimal binding to tier 2 antigen 1a123 and tier 3 antigen UKNP1.16.3 (**Fig 6G**). Neutralization potency experiments showed that class I and III bNAbs were particularly sensitive to FRLY polymorphisms at positions 431, 434, and 446 with class I and III displaying distinct neutralization profiles against tier 4 neutralization resistant strains (**Figure S10**). Further studies are necessary to uncover the contribution of each polymorphism to neutralization resistance by class I-III bNAbs.

Our findings highlight several key factors that must be considered when designing broadly protective HCV vaccines. HCV immunogens must be effective against circulating viral strains with E2 polymorphisms that contribute to resistance and immune evasion. Several substitutions that confer resistance to bNAbs recognizing the FRLY supersite have been identified^41,83^. While resistance mutations in the FRLY often lead to reduced E2 binding to CD81, loss of viral fitness, and potentially contribute to viral clearance^84^, some mutations confer resistance without a cost to viral fitness. Through comparative analysis of sensitive and resistant HCV isolates, Keck et al. identified the D431A FRLY substitution as potentially contributing to resistance to some FRLY bNAbs^39^. Further examination of isolates from individuals infected with genotype 1 indicated naturally arising D431A substitutions conferred resistance to FRLY bNAbs without decreasing viral fitness^71^. Our study further implicates the D431A polymorphism as hotspots for viral escape that must be considered in immunogen design. To promote the development of cross-reactive antibodies capable of neutralizing diverse HCV viral strains, HCV vaccines should focus the immune response on highly conserved epitopes and minimize the potential for viral escape. In a recent study, Wang et al. demonstrated the feasibility of this strategy in the context of sarbecoviruses by computationally designing receptor binding domain antigens that maximized sequence diversity at variable positions and preserved highly conserved amino acids crucial for bNAb binding. When displayed on mosaic nanoparticles, these antigens redirected antibody responses away from variable to conserved epitopes by preferentially stimulating B cells with cross reactive B cell receptors capable of bivalent binding to adjacent RBDs presenting conserved epitopes^85^. Our identification of three binding classes among HCV bNAbs targeting the key neutralizing epitopes in E2 FRLY provides a rational for immunogen design efforts to focus the B-cell response on the most conserved residues in FRLY antigenic region.

## MATERIALS AND METHODS

### IgG Expression and Purification

Genes encoding for the variable domains of HEPC74^86^, HEPC108^75^, AR3C^61^ were synthesized as gene fragments (Twist Bioscience) and cloned into the pTT5 expression vector (NRC Biotechnology Research Institute). Appropriate heavy and light chain pairs were co-transfected into Epi293F cells (Gibco) according to manufacturer’s recommendations. IgGs were isolated from filtered culture supernatants using a HiTrap Protein A HP column (Cytiva) followed by size exclusion chromatography using a Superdex200 Increase 10/300 GL column (Cytiva). Fractions corresponding to the correct mass were pooled and storaged in 1X TBS at -80°C.

### Expression and Purification of E2ecto-HEPC108 Fab Complex for Structural Studies

Untagged E2ecto from the HCV 1b09 strain was co-transfected with 6X His-tagged HEPC108 Fab heavy and untagged light chain into Epi293F cells grown in the presence of 5 µM kifunensine. The culture supernatant was filtered and the E2ecto-HEPC108 Fab complex was isolated using a HisTrap HP column and further purified by size exclusion chromatography. Fractions corresponding to the complex were pooled and concentrated to 10 mg/mL for crystallization.

### Crystallization of E2-HEPC108 Fab Complex

Initial crystallization conditions were screened using commercially available screens (Hampton Research and Molecular Dimensions) by the vapor diffusion, sitting drop method. E2ecto- HEPC108 Fab complex crystals were grown at room temperature using 0.2 µL of protein (at 15.2 mg/mL in 1X TBS) and 0.2 µL mother liquor (0.2 M potassium sodium tartrate tetrahydrate pH 7.4, 20% (w/v) PEG 3,330), cryoprotected with mother liquor supplemented with 25% (w/v) glycerol, and flash frozen in liquid nitrogen. Data were collected from a single crystal *via* fine-phi slicing using 0.2° oscillations at the Stanford Synchrotron Radiation Lightsource on beamline 12-2 using a PILATUS 6M detector. X-ray diffraction data were collected to 2.68 Å.

### Processing and Refinement of Crystallographic Data

Crystallographic data were processed using iMosfilm^87^, scaled using Aimless (CCP4)^88^, and solved by molecular replacement in Phaser (Phenix)^89^ using separate search models of 1b09 E2ecto (PDB: 6MEI) and Fab HEPC74 (PDB: 6MEH). To prepare the Fab model, SCULPTOR (CCP4i2)^90^ was used to trim side chains from regions with poor sequence similarity and the CDRH3 loop was deleted from the model. Molecular replacement successfully placed four E2ecto-HECP108 Fab complexes in the asymmetric unit. The model was refined and validated using Phenix.refine^91^ with TLS parameters and Ramachandran restraints. Iterative manual model building was performed using Coot.^92^ Glycans were initially inferred and modeled using F_O_-F_C_ maps calculated with model phases contoured to 2σ. Proper glycan geometry and restraints were generated using Privateer (CCP4i2)^93^. The structure refined to a final R_factor_ and R_free_ of 21.1% and 26.9%, respectively and **Table S1** shows data processing and model refinement statistics.

The final model quality was examined using MolProbity^94^. The model was superimposed onto previously published structures and figures rendered using PyMOL (Version 2.5.4, Schrödinger, LLC). Buried Surface Area (BSA) was calculated using the PDBePISA web tool^95^ with potential hydrogen bonds considered at a distance of < 4.0 Å and an A-D-H angle of > 90° and a maximum distance of van der Waals interaction of 4.0 Å. Rmsd calculations of model Cα alignments were performed in PyMOL without excluding outliers.

### ELISA Binding Assays

E2 ectodomain proteins were coated onto 384-well plates (ThermoFisher) at 1 µg/mL diluted in 1X TBS overnight at 4°C. Plates were washed once with 1X TBST buffer (TSB with 0.05% Tween-20) using a BioTek 405 TS microplate washer (Agilent). Plates were then blocked in blocking buffer (1X TBST, 1 % goat serum, 1 % powered nonfat milk) for 1 hr. Following blocking, plates were washed twice with 1X TBST buffer. Purified IgGs were assayed in quadruplicate with 5-fold serial dilutions starting at 10 µg/mL diluted in blocking buffer and incubated for 1 hr. Goat Anti-Human IgG HRP (Southern Biotech) was added to plates at a 1:5,000 dilution and incubated for 30 min. Plates were washed four times with 1X TBST buffer and developed upon addition of 1-Step Ultra TMB-ELISA substrate (ThermoFisher) then quenched with 1M HCl. The optical density was then read at 450 nm using a BioTek Epoch Microplate Spectrophotometer (Agilent). A non-linear regression analysis was performed using Prism version 9 (GraphPad) to calculate EC_50_ values and areas under the curve.

### Molecular Dynamics Simulations

All-atom MD simulations of the HCV E2 ectodomain (excluding HVR1, residues 414-646) were performed using GROMACS^96^ version 2024.2 using CHARMM36^97^ (July 2022) forcefield. The initial configuration of the HCV E2 ectodomain was obtained from the structure of the E2- HEPC108 complex and glycans were removed. This structure was centered in a dodecahedral simulation box with a minimum distance of 1 nm between the protein and the box edge. The system was solvated with 11,746 TIP3P water molecules, and counterions (Na^+^, Cl^-^) were added to neutralize the system and to maintain a salt concentration of 0.1 M. Energy minimization of the system was performed using the steepest descent algorithm for up to 50,000 steps, with a force convergence criterion of Fmax < 100 kJmol^-1^nm^-1^. This was followed by equilibration for 1 ns under the NVT ensemble at 310 K with position restraints on protein heavy atoms, and 100 ns in the NPT ensemble at the same temperature and 1 bar pressure, to stabilize the system’s density. Temperature was maintained using the velocity rescale thermostat with a coupling constant of 1 ps, and pressure was maintained using the C-rescale barostat with a relaxation time of 2 ps and an isothermal compressibility of 4.5 × 10^-5^ bar^-1^. Long-range electrostatic interactions were handled using the Particle Mesh Ewald method. A simulation time step of 2 fs was used, with periodic boundary conditions. Two independent 5 µs production runs were performed (total 10 µs) which were used for analysis.

Accessible surface area (ASA) of residues Y613 and W616 was calculated using the GROMACS ‘sasa’ module with the default solvent probe radius of 0.14 nm. Binding energies of the E2- antibody interactions for FRLY antibodies HEPC74 (class I), HEPC108 (class II), and AR3C (class III), were estimated computationally. Each antigen-antibody complex was simulated for 1.5 µs across three independent 500 ns runs, using the same equilibration protocol as for the E2 alone. The starting configurations for the E2-HEPC74 and E2-AR3C complexes were sourced from the PDB with IDs 6MEH and 4MWF, respectively. Missing E2 residues were not modeled. Binding energies were calculated using the molecular mechanics Poisson–Boltzmann surface area (MM-PBSA^98^) method, implemented in gmx_MMPBSA^99^ with default parameters.

### Metadynamics Simulations

Enhanced conformational sampling along the order parameter θ_*α*1-helix_ was performed using a 500 ns long well-tempered metadynamics (wt-metad) simulations^69^ implemented in GROMACS 2023.5 package patched with PLUMED-2.9.1. In wt-metad, a history dependent bias potential V(θ_*α*1-helix_, t) (defined in Eq. 1) is added to force the system overcome the kinetic barriers along the biased variable.

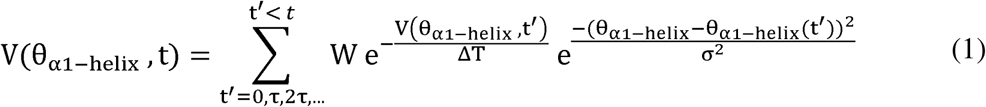

Here, τ is time period for addition of Gaussian bias (τ = 1 ps), W is the initial height of Gaussian bias (W = 1.2 kJ mol^-1^), ΔT relates to bias factor (γ) of wt-metad as γ = (T + ΔT)/ΔT, where T is simulation temperature (γ = 10), σ is the width of Gaussian bias (σ = 0.08 radian). Free energy profile along θ_*α*1-helix_ at t_end_= 500 ns was computed as F(θ_*α*1-helix_) = −γ V(θ_αl-helix_, t_end_).

### Hepatitis C Virus Pseudoparticle (HCVpp) Neutralization Assays

As previously described^100^, HIV group-specific antigen (Gag)-packaged HCV_pp_ were produced by lipofectamine-mediated transfection of HCV E1E2 plasmid, pNL4-3.Luc.R-E-plasmid bearing the *env*-defective HIV proviral genome (National Institutes of Health AIDS Reagent Program) and pAdVantage (Promega, Madison, WI) into CD81- knockout HEK293T cells.

Neutralization assays were performed as described previously^100^. HCVpp from HCV E1E2 strains UKNP5.2.1, 1a116, 1a80, and 1a142, plus strains from tier 1-4 variants (tier 1: UKNP1.11.6, tier 2: 1b34 and 1a123, tier 3: UKNP1.16.3 and UKNP1.0.1, tier 4: UKNP2.4.1 and UKNP3.1.2) derived from a panel of genetically and antigenically diverse HCVpps^76^ were used in neutralization assays. Representative mAbs from class I-III (HEPC108, HEPC74, and AR3C) were serially diluted 5-fold from a starting concentration of 10 µg/mL and incubated on Huh7 target cells for 1 hr at 37°C prior to addition to Huh7 target cells with nonspecific human IgG control. HCVpps were removed from Huh7 cells and incubated in phenol-free media for 72 hr at 37°C. HCVpp cellular entry was determined by measurement of luciferase activity in cells lysate in relative light units (RLU) as compared to PBS control. Percent neutralization was calculated as [1-(RLU_mAb_-RLU_PBS_)] x 100, with values averaged across 2 technical replicates. Log_10_ 50% inhibitory concentrations (log_10_IC_50_) were calculated from neutralization curves fit to nonlinear regression log[inhibitor] vs. normalized response, variable slope in Prism 10 software (GraphPad, San Diego CA).

## Acknowledgements

We thank members of the Flyak Laboratory for continuous support and helpful discussions, Katherine McKane and Bella Conety for help with protein expression and purification, and Benedito Melito for help with ELISA binding assays. The authors additionally thank the Cornell BioHPC facility for its computational resources. This research was supported by the U.S. National Institutes of Health (NIH) (NIH grant U19 AI159822 to A.I.F. and J.R.B.) (NIH grant R01AI127469 to J.R.B. and A.I.F) (NIH grant T32 AI145821 to X.E.W) (content is solely the responsibility of the authors and does not necessarily represent the official views of the NIH). Use of the Stanford Synchrotron Radiation Lightsource, SLAC National Accelerator Laboratory, is supported by the U.S. Department of Energy, Office of Science, Office of Basic Energy Sciences (contract no. DE-AC02-76SF00515). The SSRL Structural Molecular Biology Program is supported by the DOE Office of Biological and Environmental Research and by NIHGMS (P41GM103393).

## Data Availability

Atomic coordinates and structure factors have been deposited in the Protein Data Bank under the accession number PDB ID: 9O3D

## Author Contributions

X.E.W. and A.I.F conceived of the project. X.E.W performed experiments, processed and refined crystallographic data, and analyses. R.P. performed molecular dynamics simulations and analyses. J.M. performed neutralization experiments. N.F. performed global substitution prevalence analyses. X.E.W, R.P., and A.I.F wrote the original draft of the manuscript. All authors edited the manuscript and support its conclusions. J.R.B. and A.I.F supervised the research.

---

**Figure S1.**
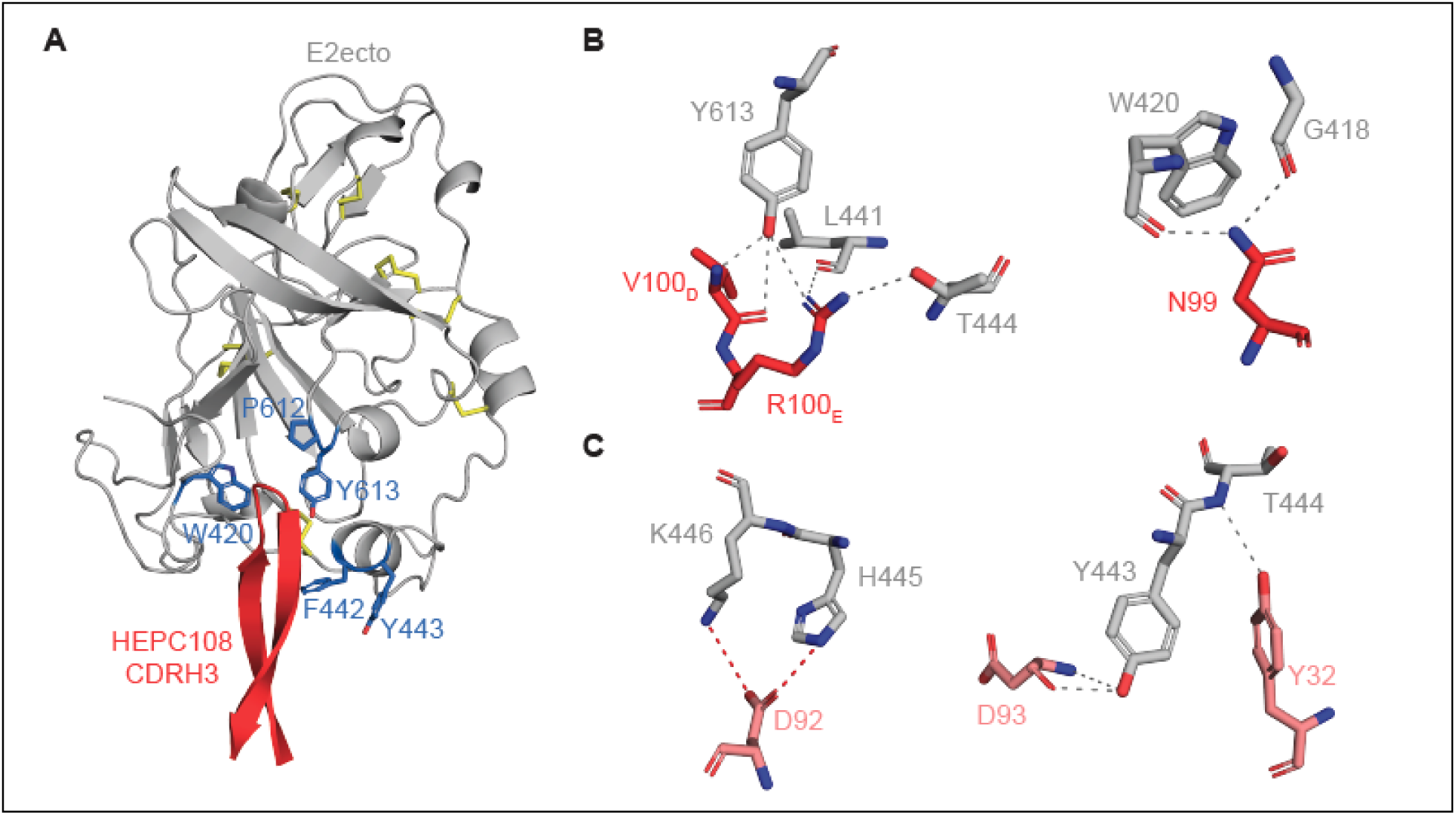
Key interactions between class II bNAb HEPC108 and the E2 FRLY. **(A)** E2ecto in complex with HEPC108 Fab showing the CDRH3 loop, colored in red cartoon, interacting with an E2 hydrophobic pocket comprised of W420, P612, Y613, F442, Y443, shown in blue sticks. E2ecto is colored in grey with disulfides shown in yellow sticks. **(B)** Hydrogen bonds between E2ecto residues and HEPC108 HC and **(C)** LC shown as grey dashes and salt bridges shown in red dashes.

**Figure S2.**
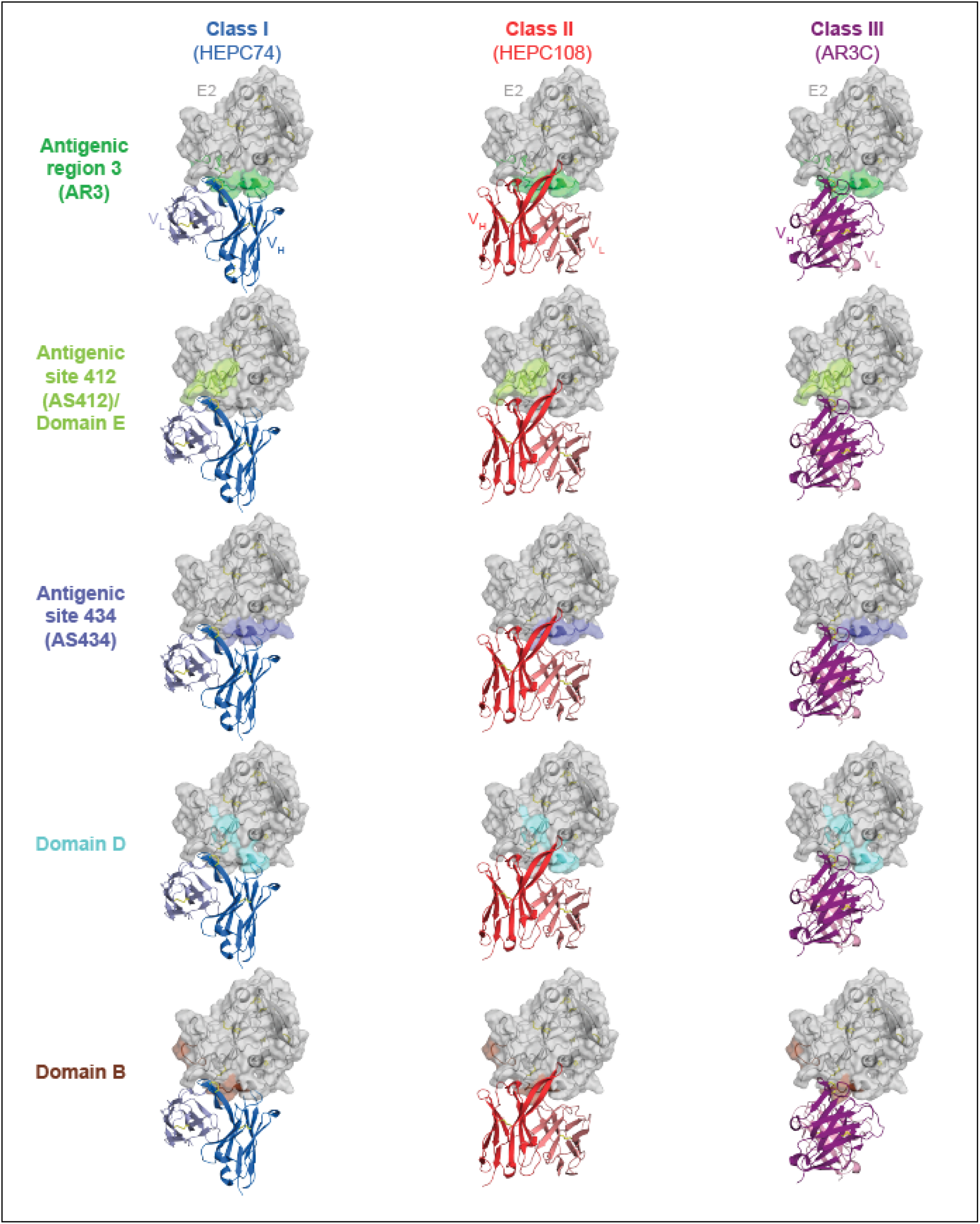
Mapping of class I-III bNAbs on previously described E2 FRLY antigenic regions, antigenic sites, and antigenic domains. Variable heavy (VH) and variable light (VL) domains of representative bNAbs HEPC74 (class I), HEPC108 (class III) and AR3C (class III), colored as in Figure 2 mapped onto E2 surface (grey surface and cartoon) showing antigenic region 3 (AR3; residues 426-443, 529-531, colored in green), antigenic site 412 (AS412; residues 412-423, colored in light green), antigenic site 434 (AS434; residues 434-446, colored in blue), domain D (residues 431-439, 529-535, colored in light blue), and domain B (residues 420-428, 441-443, 616, colored in brown). E2 and Fab disulfides show in yellow sticks.

**Figure S3.**
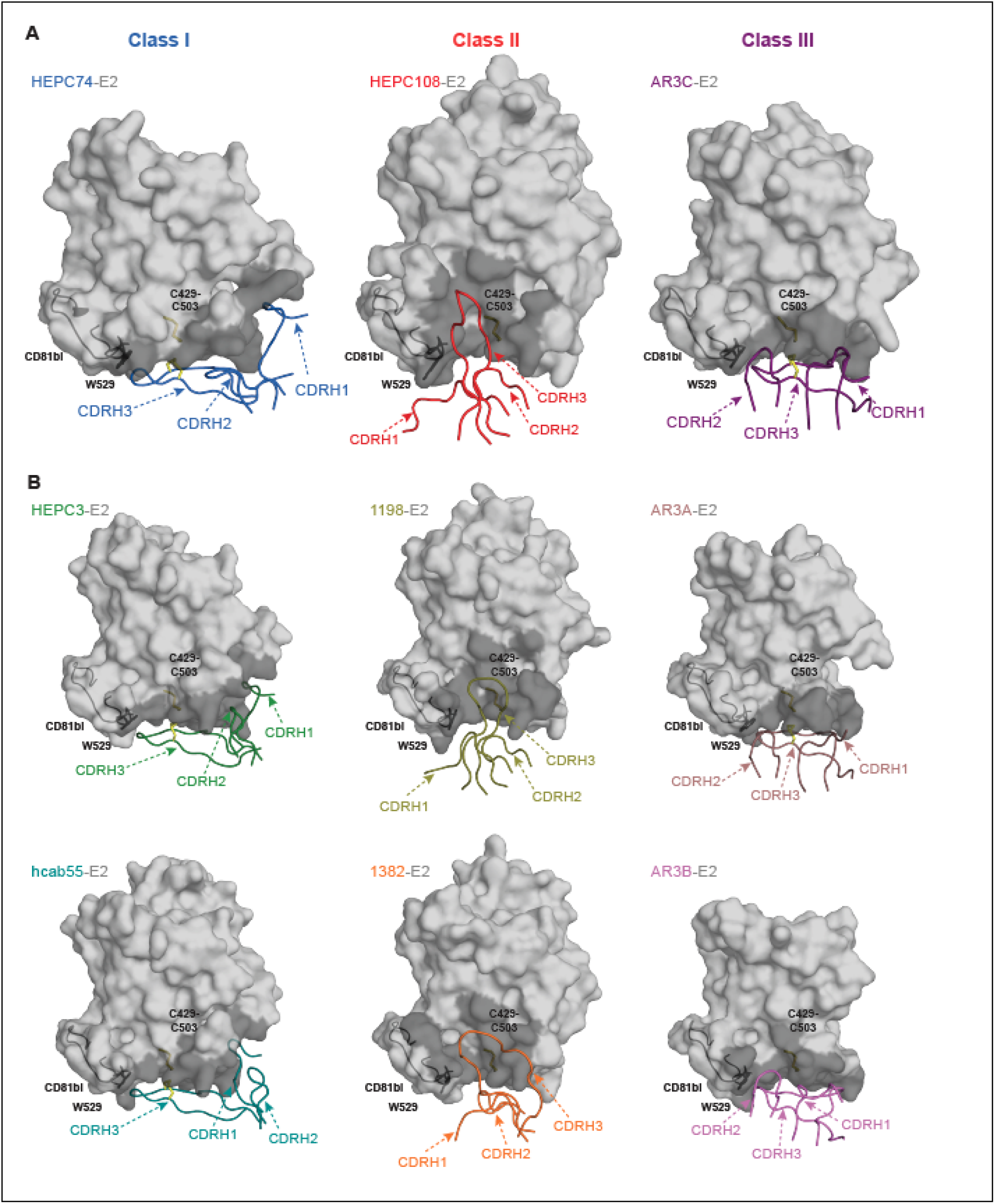
CDRH footprints of E2 FRLY class I-III bNAbs. Heavy-chain CDR (CDRH) loops for **(A)** representative bNAbs HEPC74 (class I), HEPC108 (class II), and AR3C (class III) and **(B)** additional class I (HEPC3, hcab55), class II (1198, 1382), and class III (AR3A, AR3B) members mapped onto the E2 surface. CDRH loops are colored as in Figure 2 with epitopes depicted in dark grey. The E2 CD81 binding loop (CD81bl) with conserved residue W529 are shown as black cartoons. E2 C429-C503 and Fab CDRH disulfides are shown in yellow sticks.

**Figure S4.**
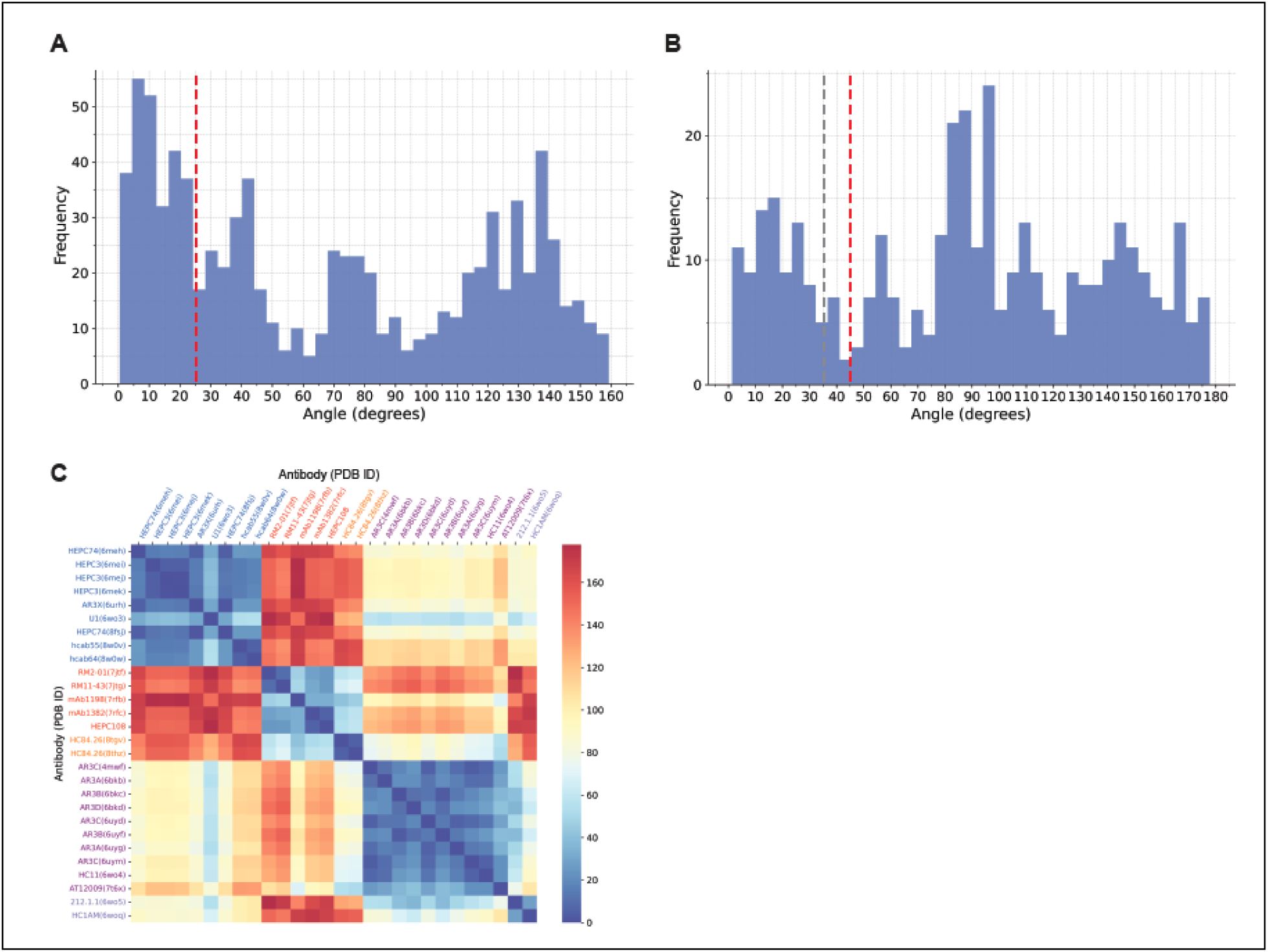
Cutoff selection for antibody binding mode clustering. Histogram of **(A)** inclination angles (φ) calculated for all antibody pairs and **(B)** approach angles (ψ) calculated specifically for FRLY-binding antibodies. The red dashed lines indicate the cutoff values selected for the DBSCAN cluserting algorithm, approximately corresponding to the first local minima in the respective histograms. The gray dashed line in **(B)** indicates the cutoff angle used for clustering in **(C)**.

**Figure S5.**
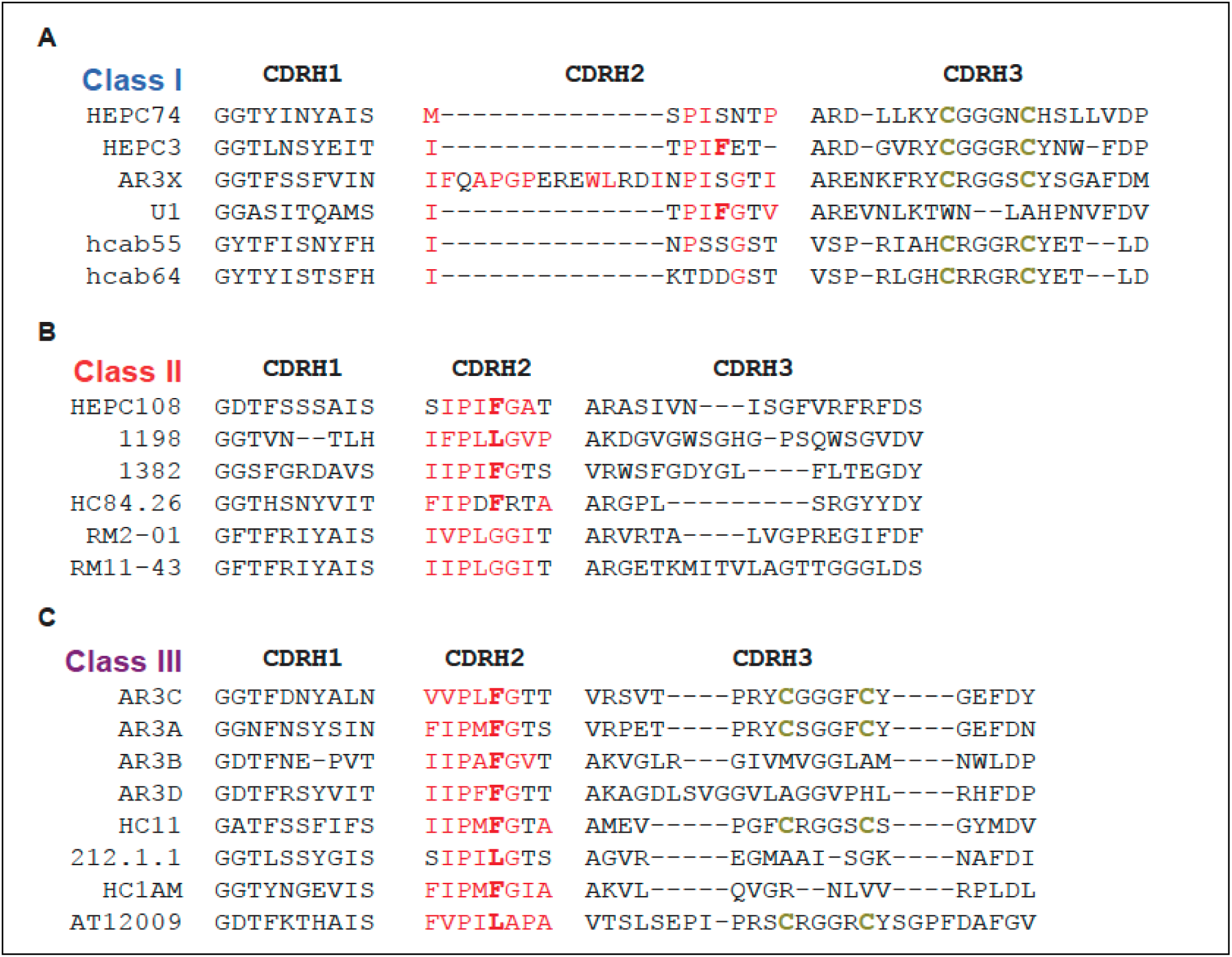
Unique CDRH sequence features of Class I-III bNAbs. Multiple sequence alignment of FRLY class I-III bNAbs. **(A)** class I (HEPC74, HEPC3, AR3X, U1, hcab55, hcab64), **(B)** class II (HEPC108, 1198, 1382, HC84.26, RM2-01, RM11-43), and **(C)** class III (AR3C, AR3A, AR3B, AR3D, HC11, 212.1.1, HC1AM, AT12009). CDRH loops were defined according to IMGT nomenclature. Hydrophobic residues in CDRH2 are colored in red, with F/L54 bolded, and cysteines within the CDRH3 disulfide motif are colored bolded gold. Dashes indicate gaps.

**Fig S6.**
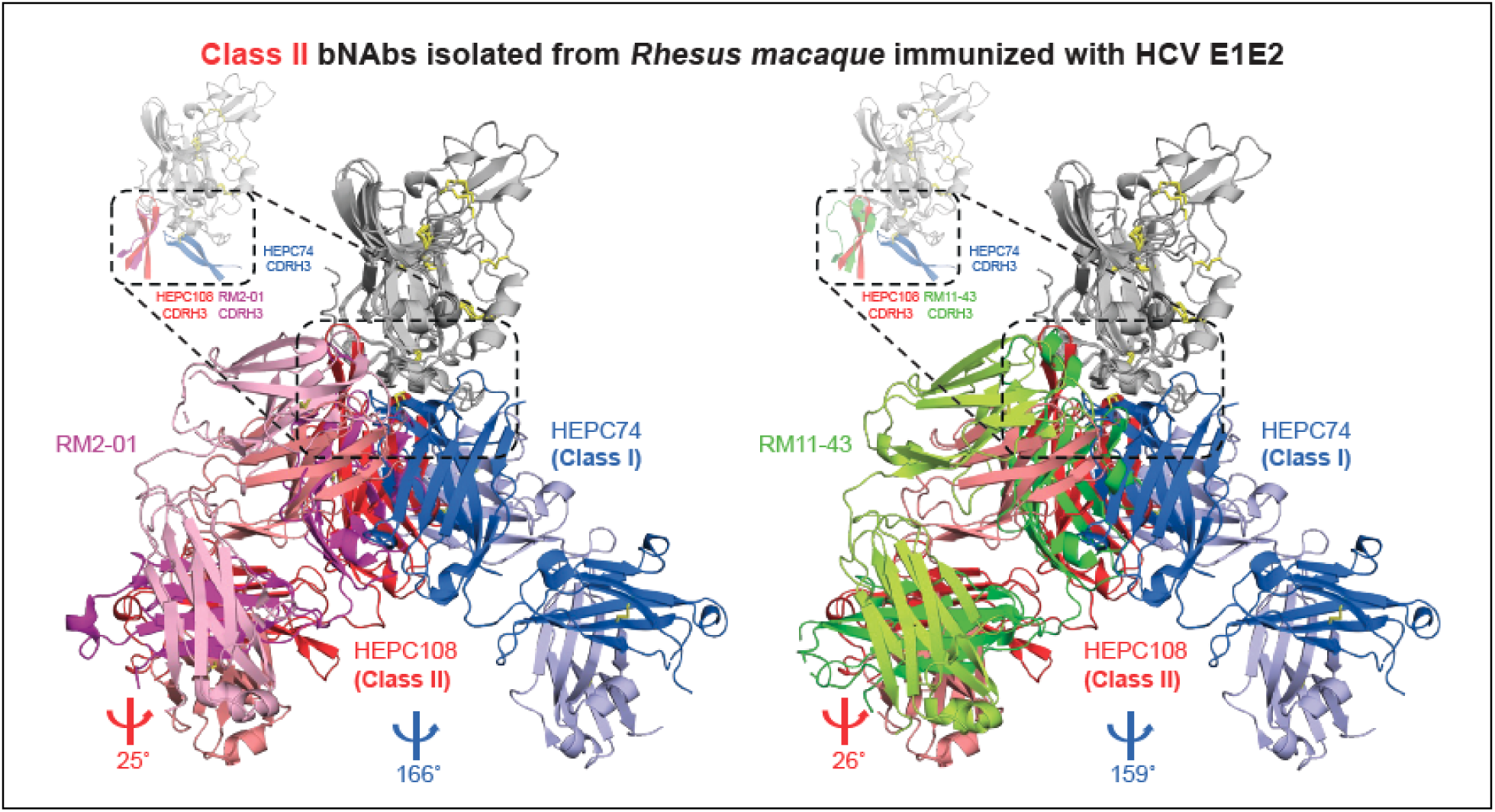
E2 FRLY class II bNAbs can be elicited by E1E2 vaccination in non-human primates. Structures of E2core in complex with RM2-01 (PDB ID: 7JTF) and RM11-43 (PDB ID: 7JTG) Fabs isolated from E1E2 immunized non-human primates aligned with class I (HEPC74)- and class II (HEPC108)-E2ecto complexes on the antigen. Coloring of HEPC74 and HEPC108 is consistent with **Figure 1**. RM2-01 HC and LC are colored in magenta and light pink, respectively. RM11-43 HC and LC are colored in green and light green, respectively. Dashed boxes denotes the E2 FRLY and CDR loops of the Fab. Left inlay displays the E2 FRLY and the Fab CDRH3 loops, highlighted within a dashed box. Red and blue angle symbols denote antibody approach angles.

**Figure S7.**
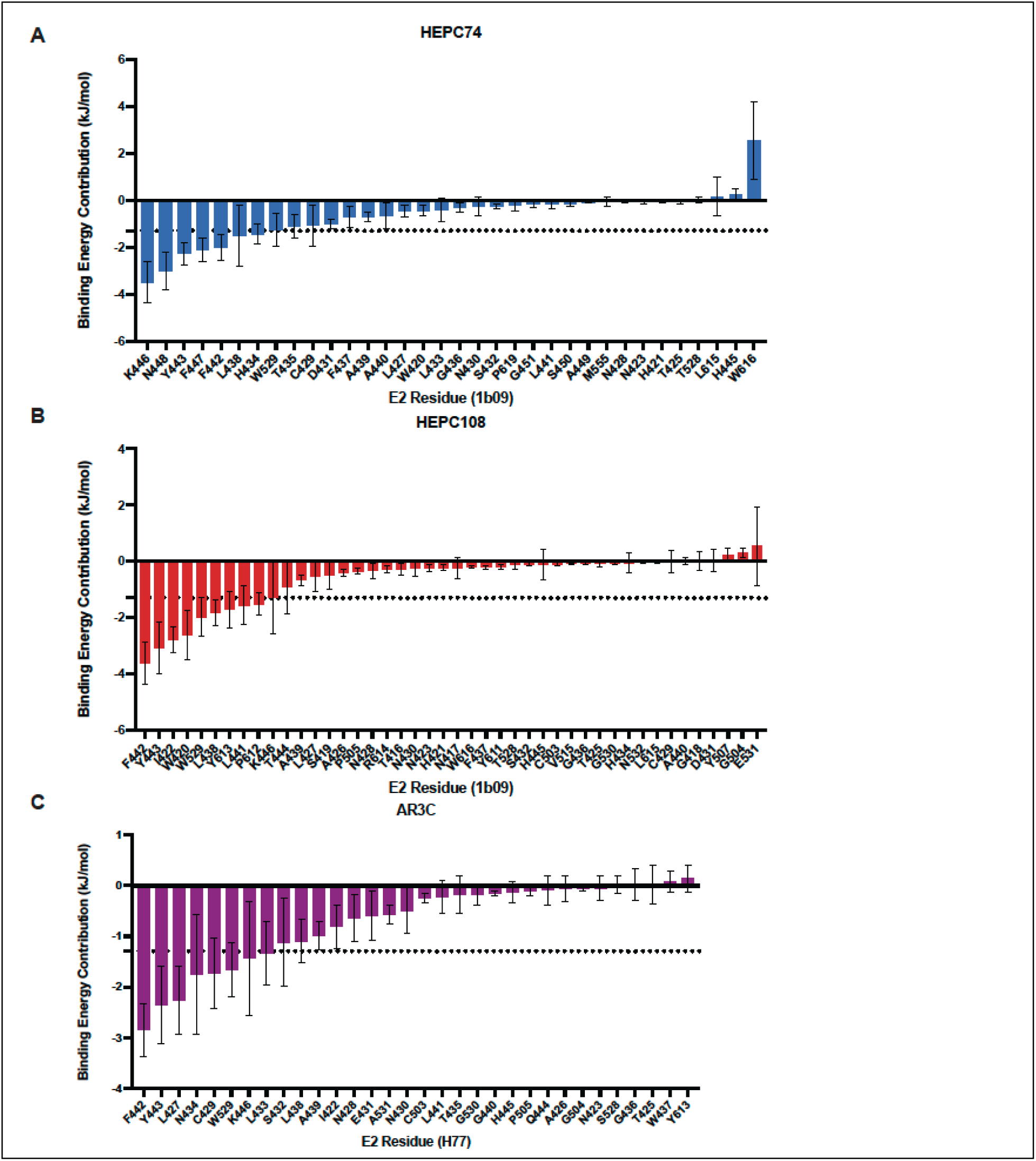
Residue-wise binding energy decomposition of HCV E2 with FRLY-specific antibodies. Per-residue contributions of HCV E2 residues to binding affinity for **(A)** E2-HEPC74 (class I), **(B)** E2-HEPC108 (class II), and **(C)** E2-AR3C (class III), calculated using MM-PBSA analysis of three independent 500 ns (total 1.5 μs) MD simulations of E2-antibody complexes. Error bars represent standard errors in the binding energy estimates. The dashed line indicates a cutoff of 1.3 kJ/mol (approx. 1/2 k_B_T,where k_B_ is the Boltzmann constant and T is the temperature), used to identify E2 residues with strong interactions with the antibody.

**Figure S8.**
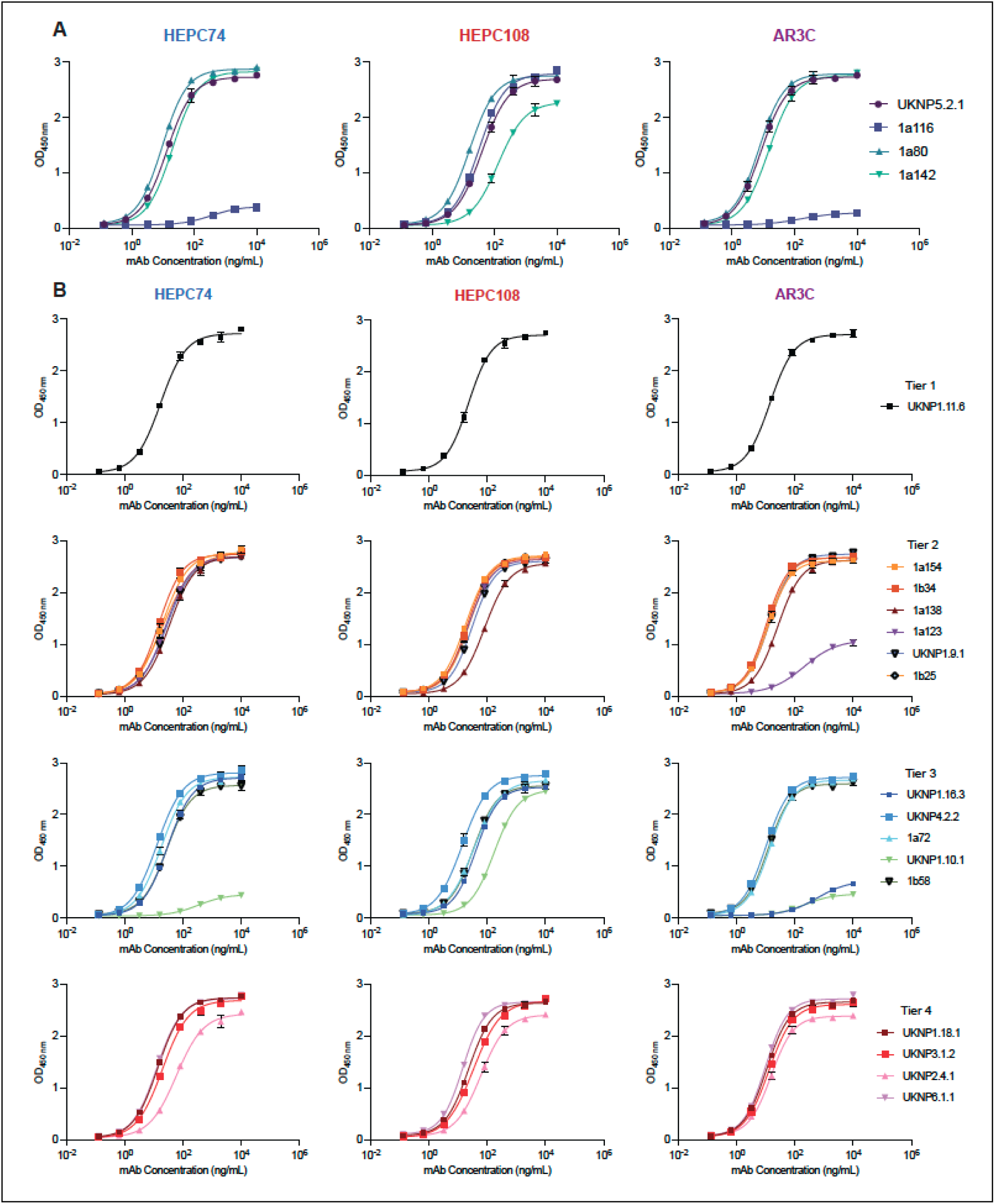
ELISA binding data of HEPC74 (class I), HEPC108 (class II), and AR3C (class III) bNAbs to E2ecto from HCV strains. **(A)** UKNP5.2.1, 1a116, 1a80 and 1a142; **(B)** Tier 1 strain UKNP1.11.6; tier 2 strains 1a154, 1b34, 1a138, 1a123, UKNP1.9.1, and 1b25; tier 3 strains UKNP1.16.3, UKNP4.2.2, 1a72, UKNP1.10.1 and 1b58; and tier 4 strains UKNP1.18.1, UKNP3.1.2, UKNP2.4.1, UKNP6.1.1. Means ± SD are shown for triplicate measurements of one representative of three independent experiments.

**Figure S9.**
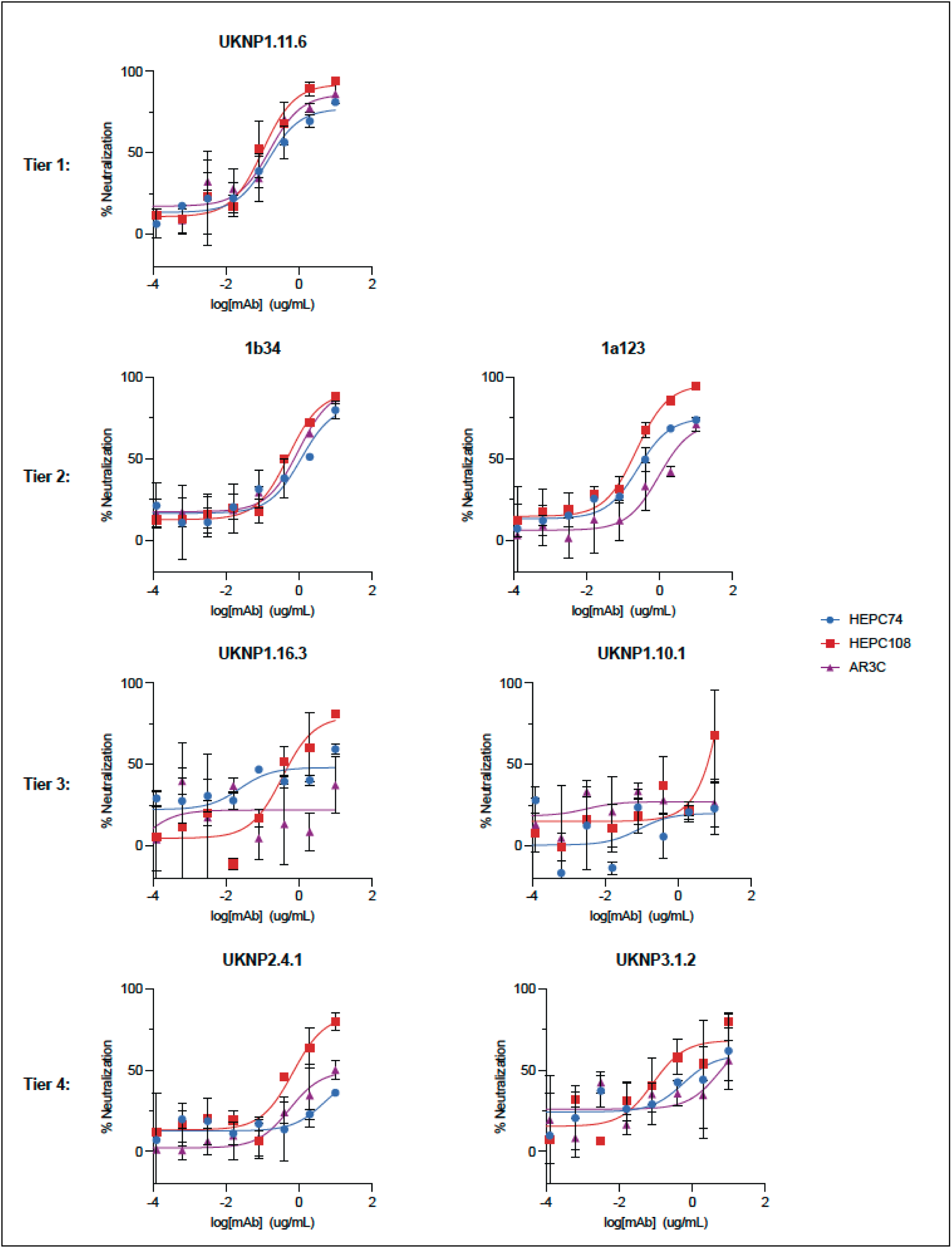
Neutralization potency of HEPC74 (class I), HEPC108 (class II) and AR3C (class III) bNAbs against. **(A)** Tier 1 strain UKNP1.11.6, **(B)** Tier 2 strains 1b34 and 1a123, **(C)** Tier 3 strains UKNP1.16.3 and UKNP1.10.1, and **(D)** Tier 4 strains UKNP2.4.1 and UKNP3.1.2. Error bars represent SD of duplicate measurements.

**Figure S10.**
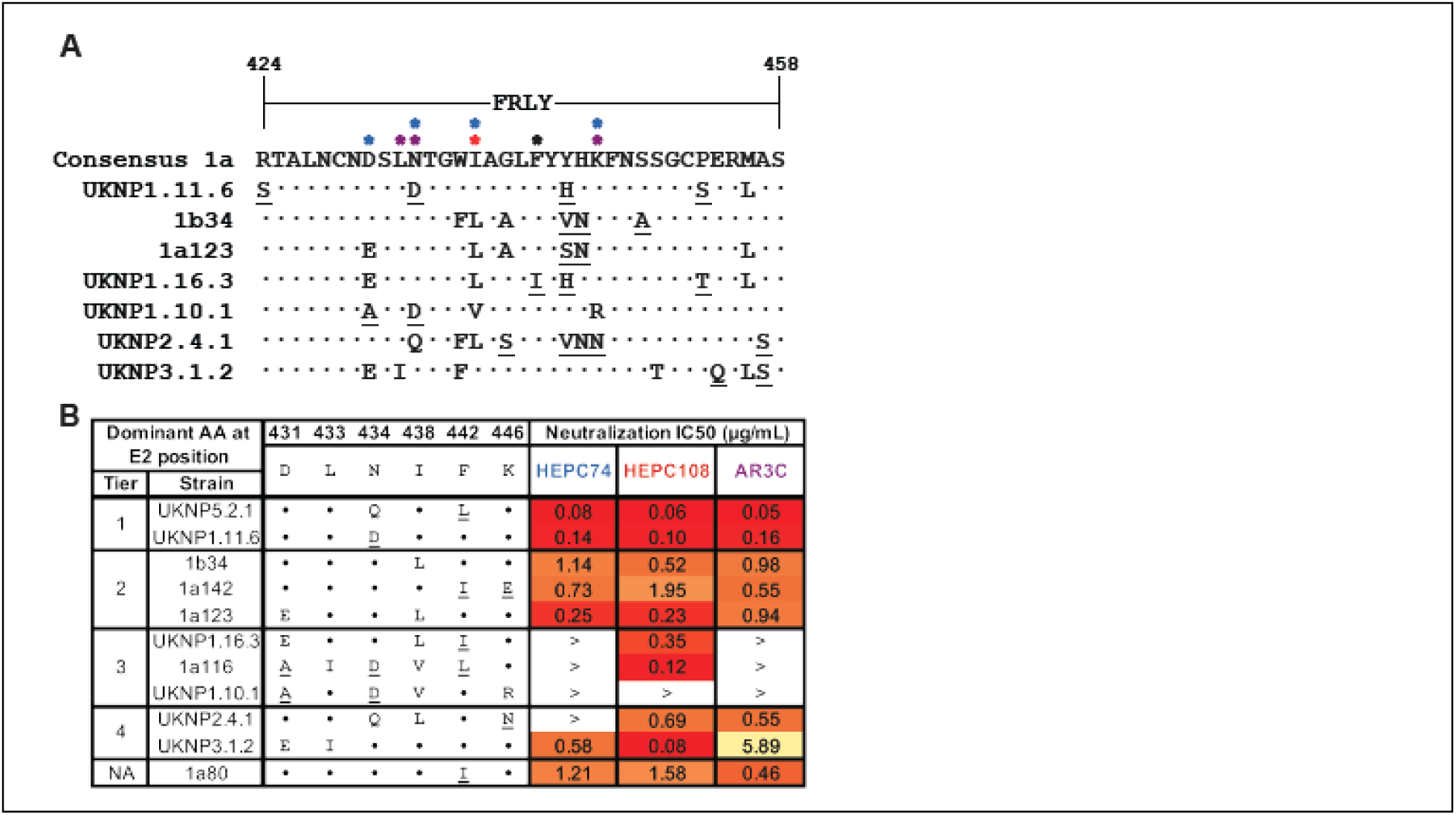
**(A)** Multiple amino acid sequence alignment of the FRLY of tier 1-4 E2 antigens tested in neuralization potency assays compared to genotype 1a consensus sequence. Residues at polymorphic positions deemed to be important for class I-III bNAb binding are denoted by a blue, red, and purple asterisks, respectively, and residues important for all three bNAb classes are denoted by a black asterisk. **(B)** Impact of E2 FRLY polymorphisms on class I-III bNAb neutralization potency. The dominant amino acid amongst genotype1a at each position are derived from **Figure 6B**. Polymorphisms that alter the biochemical properties at a given position are underlined. Positions with dots indicate identity to the dominant amino acid at the respective position. IC_50_ values are reused from **Figure 6G**.

**Table S1.** X-ray data collection and refinement statistics.

| PDB ID | HEPC108 Fab |
| --- | --- |
|  | E2ecto<sub>1b09</sub><br>9O3D |
| **Data collection<sup>a,b</sup>** |  |
| Space group | C2 |
| Cell Dimensions |  |
| *a*, *b*, *c* (Å) | 233.6, 78.3, 227.8 |
| $\alpha$, $\beta$, $\gamma$ (°) | 90, 90, 90 |
| Resolution (Å) | 82.2-2.68 (2.71-2.68) |
| R<sub>merge</sub> (%) | 13.3 (161.9) |
| R<sub>pim</sub> (%) | 8.6 (104.4) |
| CC<sub>1/2</sub> (%) | 99.6 (59.8) |
| <I/σI> | 8.9 (1.2) |
| Completeness (%) | 96.2 (97.4) |
| Redundancy | 6.3 (6.4) |
| Wilson *B*-factor (Å<sup>2</sup>) | 61.3 |
| **Refinement and Validation** |  |
| Resolution (Å) | 56.9-2.68 (2.71-2.68) |
| Unique Reflections | 112,189 (3,734) |
| Number of atoms |  |
| Protein | 20,426 |
| Ligand | 841 |
| R<sub>work</sub>/R<sub>free</sub> (%) | 21.1/26.9 |
| R.m.s. deviations |  |
| Bond lengths (Å) | 0.007 |
| Bond angles (°) | 0.89 |
| Poor rotamers (%) | 2.6 |
| Ramachandran plot |  |
| Favored (%) | 93.5 |
| Allowed (%) | 6.4 |
| Disallowed (%) | 0.08 |
| Average *B*-factor (Å<sup>2</sup>) | 89.1 |

<sup>a</sup>For each structure reported, data were derived from a single crystal.
<sup>b</sup>Numbers in parentheses correspond to the highest resolution shell

**Table S2.**
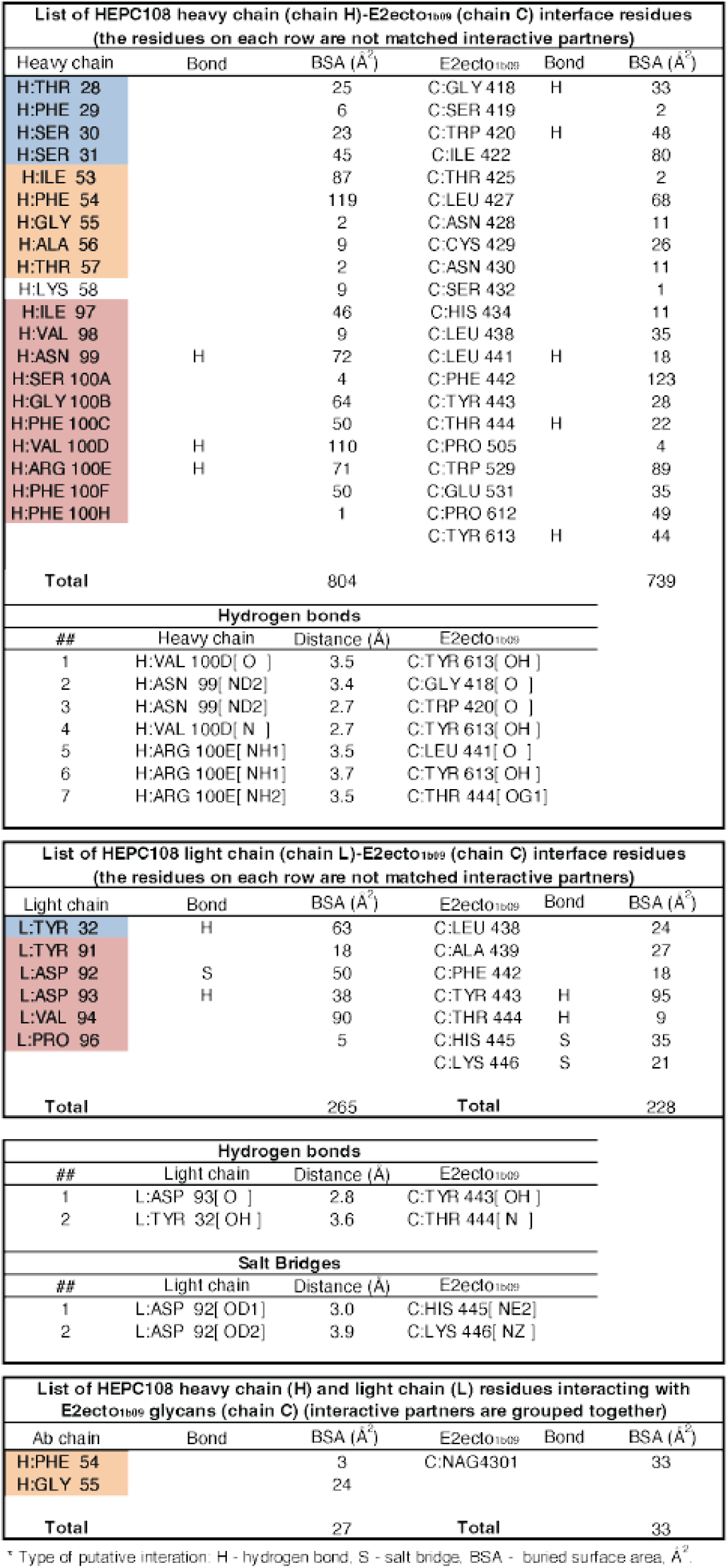
HEPC108 interface with E2ecto.

